# A comprehensive study of SARS-CoV-2 main protease (M^pro^) inhibitor-resistant mutants selected in a VSV-based system

**DOI:** 10.1101/2023.09.22.558628

**Authors:** Francesco Costacurta, Andrea Dodaro, David Bante, Helge Schöppe, Bernhard Sprenger, Seyed Arad Moghadasi, Jakob Fleischmann, Matteo Pavan, Davide Bassani, Silvia Menin, Stefanie Rauch, Laura Krismer, Anna Sauerwein, Anne Heberle, Toni Rabensteiner, Joses Ho, Reuben S. Harris, Eduard Stefan, Rainer Schneider, Teresa Kaserer, Stefano Moro, Dorothee von Laer, Emmanuel Heilmann

## Abstract

Nirmatrelvir was the first protease inhibitor (PI) specifically developed against the SARS-CoV-2 main protease (3CL^pro^/M^pro^) and licensed for clinical use. As SARS-CoV-2 continues to spread, variants resistant to nirmatrelvir and other currently available treatments are likely to arise. This study aimed to identify and characterize mutations that confer resistance to nirmatrelvir. To safely generate M^pro^ resistance mutations, we passaged a previously developed, chimeric vesicular stomatitis virus (VSV-M^pro^) with increasing, yet suboptimal concentrations of nirmatrelvir. Using Wuhan-1 and Omicron M^pro^ variants, we selected a large set of mutants. Some mutations are frequently present in GISAID, suggesting their relevance in SARS-CoV-2. The resistance phenotype of a subset of mutations was characterized against clinically available PIs (nirmatrelvir and ensitrelvir) with cell-based and biochemical assays. Moreover, we showed the putative molecular mechanism of resistance based on in silico molecular modelling. These findings have implications on the development of future generation M^pro^ inhibitors, will help to understand SARS-CoV-2 protease-inhibitor-resistance mechanisms and show the relevance of specific mutations in the clinic, thereby informing treatment decisions.

**Teaser:** Understanding how SARS-CoV-2 could counter the antiviral drug nirmatrelvir and what it means for the future of COVID-19 treatment.

## Introduction

Severe acute respiratory syndrome coronavirus 2 (SARS-CoV-2), the causative agent of COVID- 19 (coronavirus disease-19), has established itself as a permanent human and animal pathogen worldwide. Vaccines, alongside monoclonal antibodies, have drastically reduced hospitalization and/or mortality, especially in immunocompromised individuals, the elderly, and people with pre-existing medical conditions (*1*). As vaccines do not confer complete immunity against infection, the virus continues to spread effectively due to easier host-to-host transmission (*2*), and immune-escaping variants, such as Beta, Delta and Omicron (*3–5*).

In November 2021, the FDA granted emergency use authorization to Paxlovid, an orally administered medication that combines nirmatrelvir, the active component, and ritonavir, a pharmacokinetic enhancer. Nirmatrelvir is a highly potent protease inhibitor against the SARS-CoV-2 3CL^pro^/M^pro^, and is most relevant in the clinical setting, as it has been shown to decrease hospitalization, in-hospital disease progression and death (*6*). One year after the licensing of Paxlovid, another protease inhibitor, ensitrelvir, was approved in Japan through the emergency regulatory approval system under the commercial name Xocova (*7*), and recently received Fast Track designation by the FDA. Recently, leritrelvir (*8*) has been approved by the China National Medical Products Administration. Other oral M^pro^ inhibitors that have entered clinical trials include bofutrelvir (FB2001) (*9*), pomotrelvir (PBI-0451 (*10*), halted (*11*)), simnotrelvir (*12*), EDP-235 (*13*), and HS-10517 (GDDI-4405). SARS-CoV-2 continues to spread, and it is expected that the use of protease inhibitors could eventually lead to the selection of drug-resistant M^pro^ variants, with serious consequences for individuals who cannot benefit from vaccines, for example due to immune defects or pre-existing comorbidities.

To date, several studies have addressed the issue of nirmatrelvir-resistant/escaping variants using different approaches: highlighting mutational hotspots (14, 15), in silico investigation of specific mutations (16), studying resistance phenotypes (17–20), addressing which mutations are the most prone to decrease inhibitor susceptibility (21–23) or more comprehensive studies describing either mutation resistance, fitness costs, or both (24–27). Notably, a few groups have performed gain-of-function selection experiments using the Wuhan-1 strain of SARS-CoV-2. While this represents the most straightforward system to generate and study drug-resistant mutants, it requires government approval, BLS-3 facilities, and demands absolute caution to avoid biosafety breaches and potential release of these protease inhibitor-resistant variants.

Furthermore, owing to safety concerns, most of the studies that employed live SARS-CoV-2 were not performed with the Omicron variant (17–19). This precaution was taken, because protease resistant pre-Omicron variants would likely be unable to compete with the current Omicron strains, if released accidentally. In a preprint by Lan et al. (*28*), the authors validated the antiviral resistance of certain M^pro^ resistant variants in the Omicron-M^pro^ context by using SARS-CoV-2 replicons, circumventing potential safety issues for mutated transmissible variants. It is worth noting that the Omicron signature mutation (P132H) does not confer resistance to common inhibitors (*29–32*) but alters the thermal stability of the protease, as reported by Sacco et al. (*33*). This alteration, as well as other structural and biochemical features caused by the P132H substitution, could potentially affect the mechanism of resistance development and the relevance of mutations selected with Wuhan-1 M^pro^ for Omicron.

Recently, we described a BSL-2 mutation selection system based on a M^pro^-dependent chimeric vesicular stomatitis virus (VSV-3CL^pro^/VSV-M^pro^) (*34*). Replication of the chimeric VSV was effectively inhibited by nirmatrelvir. Using this system, we were able to select a panel of mutants and characterize them through computational and cellular methods. In the present study, we deepened our understanding of the SARS-CoV-2 M^pro^ resistance mutation landscape by selecting mutations in the Omicron M^pro^ and increasing the resistance phenotypes of an already resistant variant, namely VSV-L167F-M^pro^. By generating and characterizing the mechanism of Omicron- M^pro^ and L167F-derived mutations, individually or in combination, this study aims to advance the understanding of protease-inhibitor-resistant M^pro^ variants and provide valuable information for treatment decision-making.

## Results

### A safe method to select protease inhibitor resistant M^pro^ variants

In a previous study, we developed a safe alternative to using un-attenuated SARS-CoV-2 for the selection of inhibitor-resistant variants (*17*, *19*, *34*). The technology is based on a chimeric VSV- M^pro^ that encodes an artificial, non-functional polyprotein (G-M^pro^-L), and relies on M^pro^ activity for its replication (**Fig. 1A,B**). In the absence of an inhibitor, the protease processes the polyprotein, releasing G and L, and the virus can replicate. However, when a protease inhibitor is applied, the protease is inhibited, and the virus can no longer replicate.

**Fig. 1.**
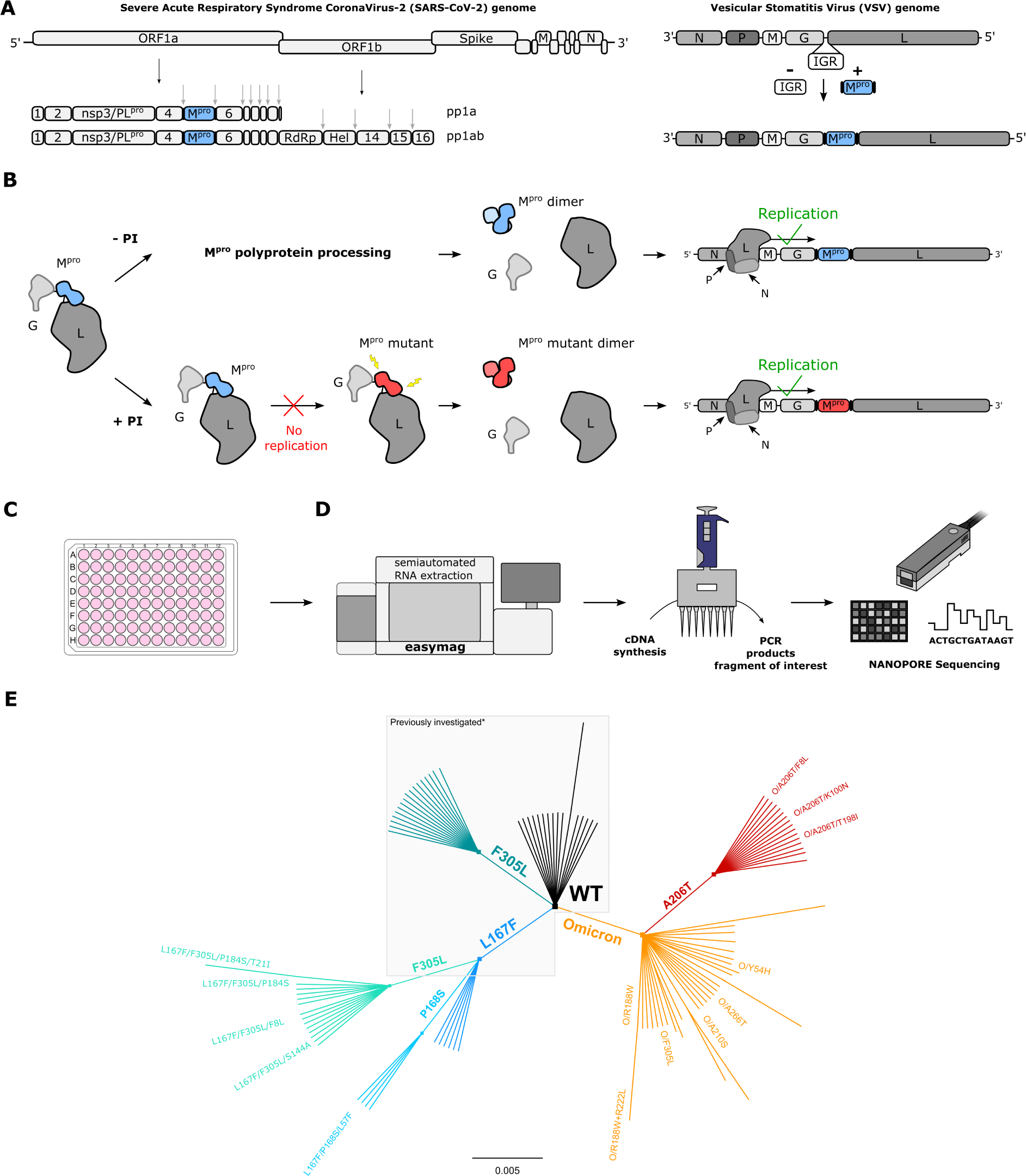
VSV-G-M^pro^-L construct: molecular mechanism, sequencing workflow, and mutant lineage phylogeny. (**A**) Schematic representation of SARS-CoV-2 genome, polyproteins pp1a and pp1ab and the VSV-based mutation selection tool. The InterGenic Region (IGR) between proteins G and L of WT VSV was replaced with the SARS-CoV-2 M^pro^ Wuhan-1 sequence and its cognate autocleavage sites. M^pro^ genome positions in SARS-CoV-2 and in VSV-M^pro^ are highlighted in light blue. (**B**) The virus is fully dependent on M^pro^ for replication. Upon translation of G-M^pro^-L, two outcomes are possible: 1. without an inhibitor, M^pro^ is free to process the polyprotein, and the transcription and replication complexes can assemble; 2. with an inhibitor, M^pro^ is inhibited, the polyprotein is not processed, and the virus is thus not able to replicate, unless it acquires a mutation rendering the M^pro^ less susceptible to the inhibitor. Then, the virus can replicate despite the inhibitor. (**C**) BHK21 cells in a 96-well plate are infected with VSV-M^pro^. Here, the two outcomes described in panel b can occur. (**D**) Nanopore sequencing workflow. (**E**) Unrooted phylogenetic tree showing the relationship between original/parental viruses and mutants. M^pro^ variants belonging to the same parental virus are coloured accordingly: WT (black), F305L (sea green), L167F (blue), L167F/F305L (light sea green), L167F/P168S (light blue), Omicron (orange), Omicron/A206T (red). For clarity, only the names of the mutants investigated in this work are displayed. *Previously generated/investigated set of mutants in our first study (*34*).

### Wuhan-1 and Omicron VSV-M^pro^ variants are equally susceptible to nirmatrelvir

To gain a deeper understanding of the potential of the Omicron-M^pro^ to acquire resistance mutations, we introduced the Omicron-M^pro^ signature mutation P132H into chimeric VSV-M^pro^, generating VSV-Omicron-M^pro^ (VSV-O-M^pro^) for subsequent selection experiments with the protease inhibitor nirmatrelvir (**Fig. S1A**). First, nirmatrelvir dose response studies were performed with the Omicron and Wuhan-1 VSV-M^pro^ in the presence of nirmatrelvir and it was found to be equally effective against both, as described previously using WT SARS-CoV-2 (*29–32*) (**Fig. S1B,C**).

### VSV-M^pro^ supports the main protease evolution of Omicron variant and variants with existing resistance to escape nirmatrelvir

Following-up on our first study (*34*), a resistant variant of VSV-M^pro^ (VSV-L167F-M^pro^), which was previously identified in selection experiments, was used. This mutation was of particular interest, because it was also found in resistance studies with authentic SARS-CoV-2 (*17–19*). Then, selection experiments using both VSV-O-M^pro^ and VSV-L167F-M^pro^ were performed. By passaging both VSV-L167F-M^pro^ and VSV-O-M^pro^ in the presence of suboptimal concentrations of nirmatrelvir, M^pro^ mutations that were generated by the error-prone VSV polymerase were selected (*35*, *36*). Samples from the first two pilot experiments with VSV-L167F-M^pro^ and VSV- O-M^pro^ were sequenced via Sanger sequencing. Samples from subsequent selection experiments were deep-sequenced (Nanopore) (**Fig. 1C,D**) on the same target region (G_Cterm_-M^pro^-L_Nterm_) (**Table S2**).

We obtained 29 distinct non-synonymous mutations in double-, triple-, and quadruple mutated VSV-L167F-M^pro^ variants and 47 distinct non-synonymous single- and double-mutated VSV-O- M^pro^ variants, as schematically shown by the unrooted phylogenetic tree (**Fig. 1E**).

To achieve multiple-mutated viruses, the most interesting mutants were selected according to specific parameters for further selection experiments, where we increased the concentration of the inhibitor, imposing stronger selection pressure at each passage. An overview on the M^pro^-genome location of the generated mutations is displayed in **Fig. S2**. Several variants were chosen for additional passaging in the presence of the PI according to the following criteria: 1. Proximity of the amino acid substitution to the inhibitor (within 5 Å of the catalytic site or near the catalytic site, within 5 – 10 Å); 2. Prevalence in the GISAID database of specific mutations (*37–39*) (**Fig. 2A,B**), and for further analyses, substitutions whose count in the GISAID database was above 500 entries, were considered (**Fig. 2A**); 3. Frequency of a specific mutation occurring in different samples independently (**Fig. 2C**).

**Fig. 2.**
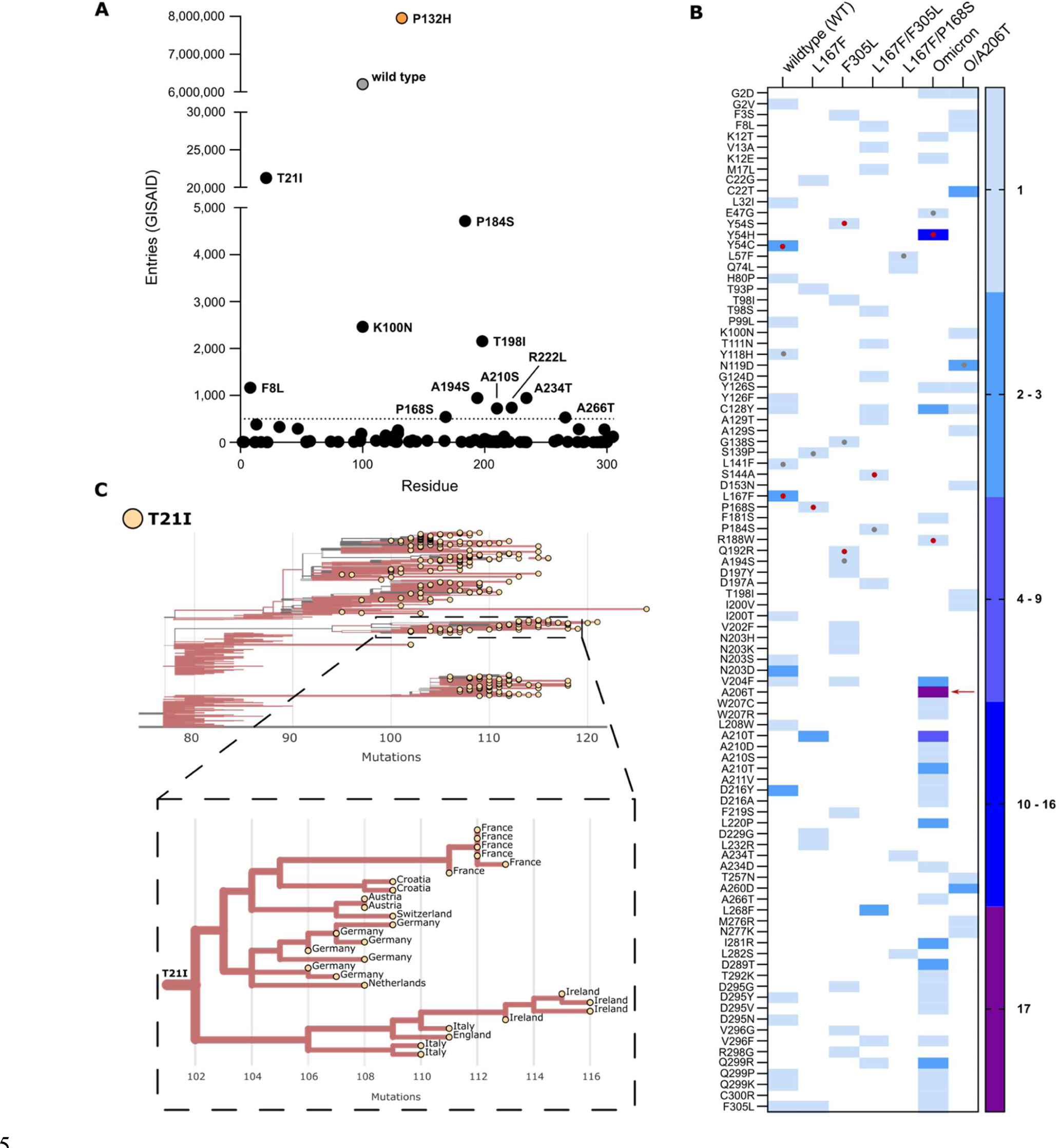
Frequency of M^pro^ mutations in the GISAID database. (**A**) The total number of nsp5/M^pro^ substitution entries for each mutation found during the selection experiments. WT and Omicron- M^pro^ sequences are displayed in grey and orange, respectively. The dotted line at 500 represent the cut-off value. (**B**) Heat map representing the number of a specific substitution’s independent occurrences from different samples of VSV-M^pro^ selection experiments. The red arrow indicates the substitution A206T. Red dots indicate residues within the catalytic site and grey dots indicate residues near the catalytic site. (**C**) Phylogenetic subtree of nsp5/M^pro^-T21I mutant generated with the Ultrafast Sample placement on Existing tRee (UShER) tool and magnified view of a subtree area, showing transmission of this variant possibly from a single founder event (9^th^ of April 2023). Only sequences deposited after the Omicron emergence were used.

Some mutations combined more than one criterion. For example, some substitutions located in the catalytic site were also frequent in the GISAID repository. However, residues located in this area were generally more conserved (**Fig. S3**) (*40*, *41*), whereas residues outside the catalytic site were more frequent in our selection experiments and more frequent in GISAID. We found ten substitutions with more than 500 entries: T21I, P184S, K100N, T198I, F8L, A234T, A194S, A210S, A266T, and P168S (**Fig. 2A**). Despite not reaching 500 entries in GISAID, we also included the F305L mutation for further study. We recently described F305L as an autocleavage site optimization mutant with similarities to T304I selected in authentic SARS-CoV-2 resistance studies. All mutations occurred during selection experiments, except for T21I and R222L. R222L arose during plaque purification of VSV-Omicron-R188W-M^pro^. T21I appeared during the generation of VSV-L167F/F305L/P184S from its plasmid (also referred to as “rescue” (*42*)). This virus variant had to be produced from its plasmid, as it could not be plaque purified.

Based on the abovementioned parameters (proximity, GISAID frequency and/or independent occurrences), two VSV-L167F-M^pro^ variants, L167F/F305L and L167F/P168S, were chosen for further selection experiments. From VSV-O-M^pro^, the variants O/A206T, O/R188W, O/Y54H, O/A210S, and O/A266T were chosen for further study. Among these variants, only O/A206T was selected for additional selection experiments. O/A206T did not fulfil the first two filtering criteria, but only the third (multiple occurrences), as it occurred 17 times independently in VSV-O-M^pro^ selection experiments. To confirm that this variant was not present in our VSV-Omicron-M^pro^ stocks already, Nanopore sequencing was performed on those stocks. We confirmed the absence of any VSV-O-M^pro^ variant subpopulation to the extent of sensitivity that Nanopore sequencing provides. Subsequently, the next round of selection experiments with the variants L167F/F305L and L167F/P168S was carried out, in which the triple-mutant variants L167F/F305L/F8L, L167F/F305L/S144A, L167F/F305L/P184S, and L167F/P168S/L57F were selected. From VSV- O/A206T-M^pro^, the mutants O/A206T/F8L, O/A206T/K100N and O/A206T/T198I were selected. To assess whether mutants generated from the selection experiments and selected for further analyses were not detrimental for viral propagation, the Ultrafast Sample placement on Existing tRee (UShER) tool was applied. Phylogenetic analyses using sequences harbouring T21I (**Fig. 2C**), T198I, P184S, K100N, and F8L (**Fig. S4A,B**) were generated. We chose these mutations from our mutation pool because they were the most frequently represented in the GISAID database. The prevalence of these mutations before and after the Omicron surge (21^st^ December 2021, up to 18th January 2023) (**Fig. S4C**) has also been examined. From this analysis we could observe that M^pro^ variants bearing these substitutions appear to not be harmful to the replication and spread of the virus.

### Mutations selected in VSV-G-M^pro^-L confer resistance to nirmatrelvir & ensitrelvir

To quantify the resistance of the selected variants, the most interesting mutations were introduced into two previously described live-cell-based protease activity assays, namely 3CL/M^pro^-On and 3CL/M^pro^-Off (*43*) (**Fig. S5B,C**). These two protease activity measurement tools rely on replication-incompetent VSV-dsRed variants missing either the phosphoprotein (P) or the polymerase (L), which are replaced with the red fluorescent protein dsRed. Briefly, cells were transfected either with a plasmid encoding the P protein modified with an INTRAmolecular-M^pro^- tag (P_Nterm_:M^pro^:P_Cterm_) or an INTERmolecular fusion protein made of the green fluorescent protein, M^pro^ and L (GFP-M^pro^-L). The cells were then infected with VSV-ΔP or VSV-ΔL, respectively. The P_Nterm_:M^pro^:P_Cterm_ intramolecular tag in combination with VSV-ΔP-dsRed constitutes a gain-of-signal assay also called “M^pro^-On”, whereas the artificial polyprotein (or fusion protein) GFP- M^pro^-L in combination with VSV-ΔL-dsRed constitutes a loss-of-signal assay also called “M^pro^- Off”.

First, the M^pro^-On system was used (**Fig. S5C**) to quantify the resistance phenotype of four triple mutants that arose from the WT protease: L167F/F305L/F8L, L167F/P168S/L57F, L167F/F305L/P184S, and L167F/F305L/S144A, against nirmatrelvir and ensitrelvir (**Fig. 3A**).

**Fig. 3.**
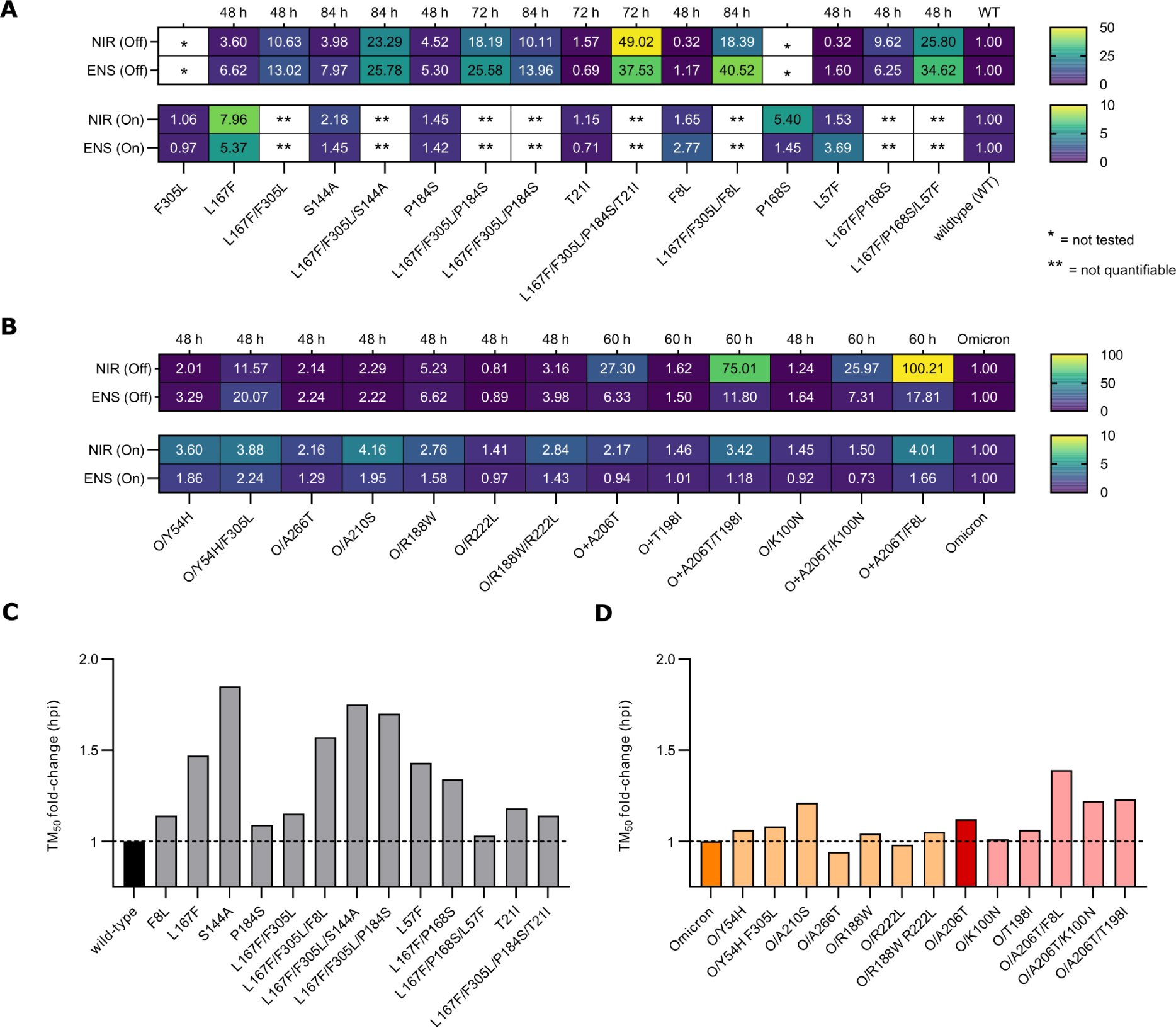
Resistance data and viral replication kinetics of WT, Omicron and mutant main proteases. Rectangular fields with one and two stars represent constructs/variants that have not been tested or for which IC50 values were not quantifiable, respectively. Hours after infection / hours post infection (hpi). (**A**) Heat maps of IC_50_ fold-changes of M^pro^-On (bottom) and M^pro^–Off (top) dose responses for nirmatrelvir and ensitrelvir. (**B**) Heat maps of IC_50_ fold-changes for Omicron-M^pro^-On (bottom) and Omicron-M^pro^–Off (top) dose responses for nirmatrelvir and ensitrelvir. (**C**) Bar plot of TM_50_ fold-changes vs the WT protease. The WT M^pro^ is coloured in black. (**D**) Bar plot of TM_50_ fold-changes vs Omicron (orange) and Omicron/A206T (bordeaux) proteases. Variants are coloured in light orange and light bordeaux according to the parental protease from which they originated.

We also included the related single and double mutants to investigate the contribution of each mutation to the resistance phenotype. Single substitutions conferred low resistance to protease inhibitors, but the resistance increased by combining them. However, there were limitations in quantifying the resistance of some WT-related combinations using the gain-of-signal assay. This was largely observed in triple mutants, where the gain in signal was strongly supressed by the high resistance of these protease variants and IC_50_ values could not be quantified. Since inhibitor concentrations of 100 µM and higher were cytotoxic, thereby interfering with signal generation, we did not increase the inhibitor concentration further. Instead, to attain better resistance quantification, we cloned the same set of WT M^pro^ mutations in the loss-of-signal assay. This cellular system is more sensitive, as the IC_50_ value of the WT M^pro-^Off is considerably lower than in M^pro^-On. Furthermore, the fluorescent signal was measured at different time points (48, 72 and 84 hours), because some variants were replicating slower, affecting final signal quantification (**Fig. 3C**). As expected, while single mutants had a mild effect on M^pro^ susceptibility to nirmatrelvir and ensitrelvir, triple mutants showed increased resistance (**Fig. S6B,D,F**). Initially, the most prominent differences between PIs were observed for L167F/F305L/F8L with 18.4-fold and 40.5- fold, and L167F/P168S/L57F, with 25.8-fold changes and 34.6-fold changes for nirmatrelvir and ensitrelvir (**Fig. 3A** and **Fig. S7**), respectively. When T21I was added to L167F/F305L/P184S, the resistance phenotype substantially increased, with 49.0- and 35.5-fold changes in the IC_50_ values of nirmatrelvir and ensitrelvir, respectively (**Fig. 3A**).

Mutations that arose during selection experiments with the Omicron protease (P132H) were also characterized in the dose-response assays. In **Fig. S8**, non-linear regression analyses of dose-response experiments are shown. IC_50_ values were extrapolated from each curve and calculated the IC_50_ fold-changes for all the variants compared to the parental protease, Omicron-M^pro^ (**Fig. 3B**). In the M^pro^-On assay, neither single nor combined mutants showed strong resistance compared to the parental protease. We therefore cloned the same set of mutations for loss-of-signal experiments to confirm the phenotype. Some Omicron-M^pro^ variants were measured at different time points to adjust for their slower kinetics. The resistance levels in the M^pro^-Off system were far greater than in the M^pro^-On assay, as noticeable from the IC_50_ fold-changes. M^pro^-O/A206T-Off, M^pro^- O/A206T/K100N-Off, M^pro^-O/A206T/T198I-Off, M^pro^-O/A206T/F8L-Off and M^pro^- O/Y54H/F305L-Off showed large differences in the susceptibility to the two inhibitors. Nirmatrelvir showed IC_50_ fold-changes of 27.3, 26.0, 75, 100.2 and 11.6, respectively. Moreover, we observed differences between responses against nirmatrelvir and ensitrelvir, in particular for M^pro^-O/Y54H/F305L-Off, which was more resistant against ensitrelvir with an IC_50_ fold-change of 20.1.

### Protease inhibitor resistant mutations can alter VSV replication kinetics, which can be restored upon acquisition of compensatory mutations

As described above, some variants, e.g. S144A and triple mutants except L167F/P168S/L57F, did not show strong resistant phenotypes at the time point we typically measure dsRed signal (48 h). To investigate this unexpected result, viral replication was assessed over time using the M^pro^-Off system. In M^pro^-Off transfected cells, the signal increases over time in the absence of an inhibitor (**Fig. S9A**). Fluorescence output was measured every 12 h, beginning 12 h after infection (= hours post infection, hpi). Fluorescent signals were then plotted against hpi and non-linear regression analyses were performed, which returned TM_50_ (time maximum fifty), a value that represents the time required to reach the half-maximum value of the curve when the signal plateaus. The WT protease had a TM_50_ between 35-40 hpi (**Fig. S9E**), and the plateau started at 48 hpi, which matched the time point that we generally consider for dose-response experiments in M^pro^-Off assays. As expected, the WT protease led to the fastest increase in fluorescence signal, and mutants showed either similar or higher TM_50_ values (**Fig. 3C, S9F,G**). This phenomenon indicates that certain resistance mutations can reduce protease activity, which is expected to attenuate virus replication if the mutations would occur in wild-type SARS-CoV-2. In contrast, by adding T21I to M^pro^- L167F-P184S-F305L-Off resulting in the quadruple-mutant M^pro^-T21I-L167F-P184S-F305L-Off, a decreased TM_50_ of 14 hours compared to its parental virus was observed (**Fig. S9H**).

We also investigated viral replication kinetics of the selected Omicron-M^pro^ mutants. Overall, mutations that have occurred in the Omicron-M^pro^ context, had a lower impact on viral replication rate than in the WT M^pro^ (**Fig. 3D**). The slowest variant was O/A206T/F8L, with a 1.4 fold-change (∼16 hours) increase in the time required to reach the plateau.

### Purification and resistance characterization of M^pro^ variants through dose response experiments via a fluorescent-based biochemical assay

To further support our findings, recombinant proteases of a few selected mutants were produced as described in **Fig. S10A** (WT, L57F, L167F/P168S and L167F/P168S/L57F) for a subsequent fluorescent-based protease activity inhibition assay. The activity of the purified proteases was tested by western blot analyses using the recombinant proteases and a cellular expressed cutting reporter bearing the nsp4/nsp5 autocleavage sequence in the SARS-CoV-2 polyprotein. The M^pro^ variants were tested also with a fluorescence cleavage-based (**Fig. 4C-E**) assay and the resistance phenotype of the L167F/P168S/L57F variant observed in our cell-based assays was confirmed. To display dose-response fitting curves from the fluorescence cleavage-based assay, we normalized to the highest average mean of each dataset independently, thereby obtaining curves with the same plot scale. As shown in **Fig. 4F**, mutant proteases had progressively decreased cleavage activity.

**Fig. 4.**
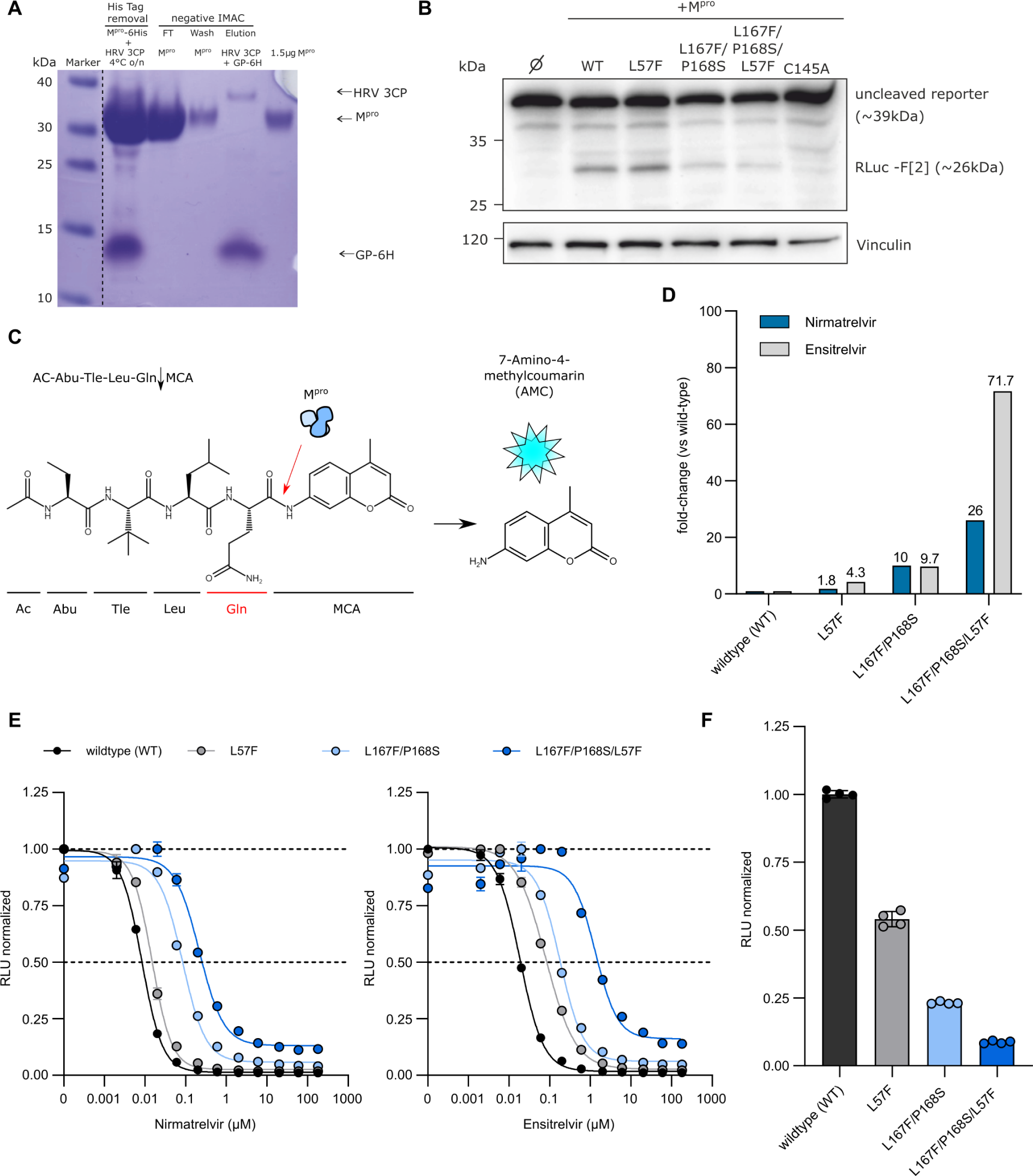
Biochemical assay dose responses of nirmatrelvir and ensitrelvir against selected mutant recombinant proteases. (**A**) Representative Coomassie-staining for the M^pro^-L57F variant after FPLC purification and His-tag removal with HRV-3CP followed by a negative Immobilized Metal Affinity Chromatography (IMAC). The dotted line indicated that the marker lane photograph was cut and placed near the production of the recombinant M^pro^ lanes. Lane 1: Tag removal reaction of M^pro^ with HRV-3CP. Combined band of M^pro^ and HRV 3CP at 35 kDa and free 6xHis-tag (GP- 6H) at around 10 kDa. Lane 2: Flow-through (FT) of negative IMAC, yields the M^pro^-L57F without His tag. Lane 3: Wash step, indicating a slight loss of protein. Lane 4: The elution step leads to the release of the bound HRV-3CP protease (His tagged) and cleaved His tag of M^pro^. Lane 5: 1.5 µg M^pro^-L57F without His Tag after desalting. (**B**) Western blot analysis of the cleavage of the “long” cutting reporter after the addition of the different M^pro^ variants. The RLuc-F[2] band at approximately 26 kDa indicates successful reporter cleavage. One representative western blot of n = 3 independent experiments is shown. (**C**) Schematic representation of the fluorescence cleavage-based M^pro^ activity assay. (**D**) Fold-changes of IC50 values of nirmatrelvir (blue) and ensitrelvir (light grey) against WT, L57F, L167F/P168S and L167F/P168S/L57F proteases. (**E**) Dose response experiment with nirmatrelvir (left) and ensitrelvir (right) with WT, L57F, L167F/P168S and L167F/P168S/L57F (± SEM; n = 2). (**F**) WT and mutant proteases relative activity at 4 hours (± SD; n = 4). The WT signal in the absence of inhibitor was normalized to 1 and mutants’ relative activities were normalized accordingly using WT as reference scale.

### Computational analyses reveal destabilization of M^pro^-nirmatrelvir complex formation and provide insights on protease stability and dimerization affinity upon mutation

To provide a molecular interpretation of the mechanism of resistant M^pro^ variants against nirmatrelvir compared to the WT, we performed molecular modelling with Bioluminate (*44–47*). This software uses already existing structural data to calculate the impact of a mutation on the stability of a specific protein conformation. The software returned models of mutant structures and delta stability values (Δ_Stability in kcal/mol), indicating whether a mutation stabilizes (negative values) or destabilizes (positive values) the investigated conformation. Many of the investigated mutations returned positive values, indicating destabilization of the nirmatrelvir-binding conformation in the WT and Omicron-M^pro^ structures, (PDB entries 8DZ2 (*48*) and 7TLL (*49*), plus related variants (**Fig. S13**)). Most notably, Δ_Stability values of +36.8, +36.5, +46.9 and +14.2 kcal/mol were calculated for the M^pro^ L167F/P168S/L57F, L167F/F305L/P184S/T21I, Omicron/A206T and Omicron/A206T/K100N mutations respectively, which were among the mutations that conferred the strongest resistance phenotypes (**Fig. 3A**, **Fig. 3B**).

Some mutated residue side chains are within the ligand binding site, therefore close to the inhibitor, but do not directly interact with it, namely L167, P168, Y54, L141 and several others (**Fig. 5A**). Furthermore, the majority of selected and observed resistance mutations were located outside of the drug-binding site. We therefore assumed that both near-catalytic site and distant mutations exert an allosteric effect. I.e., they change protein dynamics in favor of conformations that are less compatible with ligand binding, especially when in combination with other mutations.

**Fig. 5.**
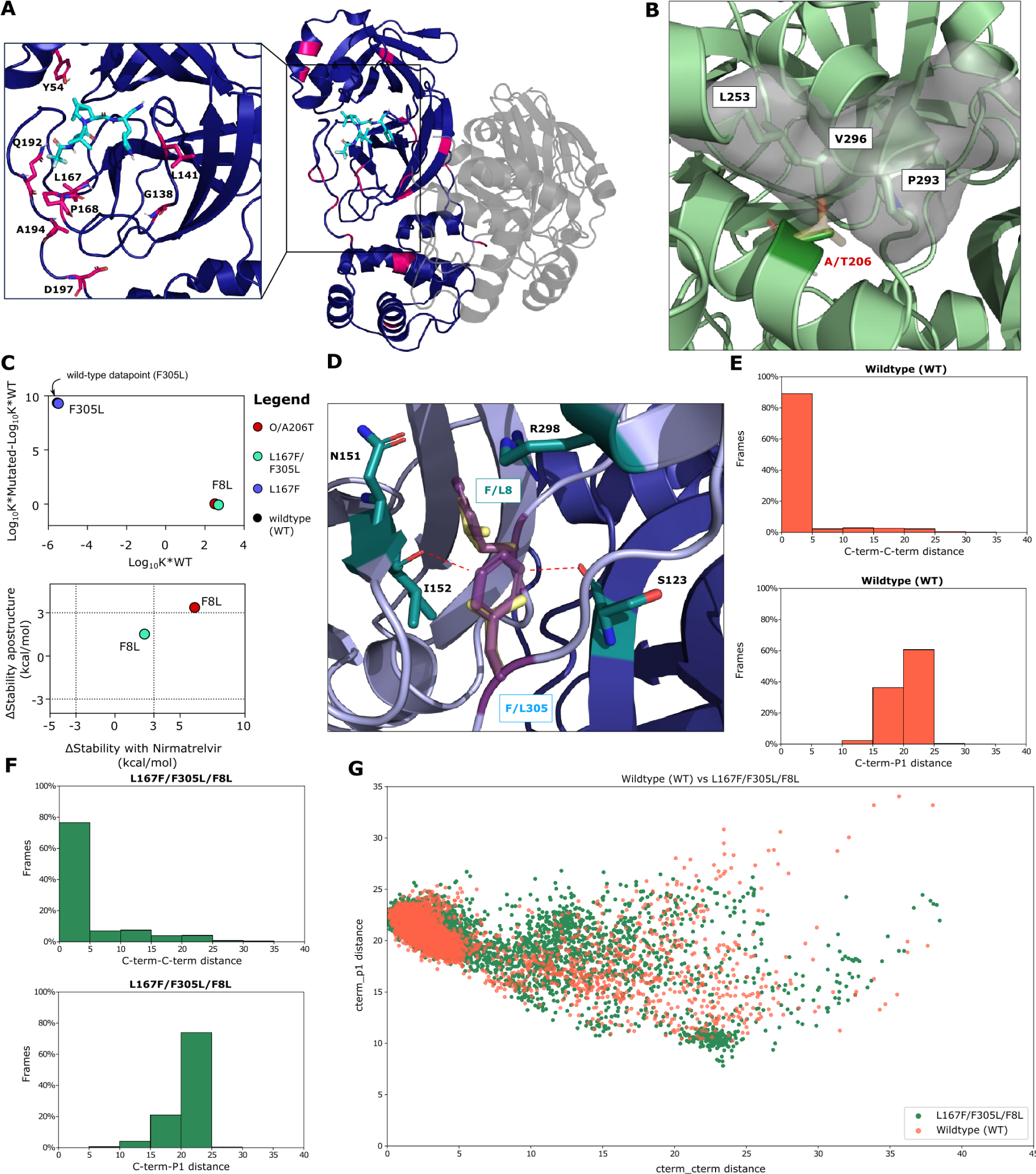
Structural analysis of mutants. (**A**) Overview of investigated mutants (pink) mapped onto the M^pro^ homodimer (dark violet and grey) bound to nirmatrelvir (light blue sticks, PDB 8DZ2) (*48*). Side chains of residues within the drug binding site are shown as sticks to highlight their distance to nirmatrelvir. (**B**) The WT alanine side chain of residue 206 (dark green sticks) is tightly packed against the hydrophobic side chains of residues L253, P293, and V296 (green sticks, PDB entry 7ALI) (*51*). The threonine side chain (yellow sticks) is polar and would clash with these residues as indicated by the light grey surface. (**C**) Dimerization affinity plot of F8L and F305L mutants (top) and Δ_Stability values (bottom) of apo and nirmatrelvir bound structures for the F8L mutants within different variant backgrounds. The additional dotted lines at x = ± 3 and y = ± 3 in the stability plot are cut-off values that we considered as a meaningful difference in protease stability (a positive value indicates a decrease in protease stability and a negative value indicates an increase in protein stability). (**D**) The F8 and F305 aromatic side chains (dark violet) form a pi- pi T-stack and F8 in addition interacts with N151, I152, and R298 (green sticks, PDB entry 7ALI (*51*)). F8 is located within the homodimer interface, with chains A and B colored light violet and blue, respectively. The conjugated p-orbitals of the F305 side chain are located nearby of the I152 and S123 backbone oxygen atoms, potentially leading to electronic repulsion. In the F305L mutant, the distance of the leucine side chain (yellow sticks) to these oxygen atoms is considerably increased. (**E**) Bar plots showing the percentage of frames against C-term/C-term distance or C- term/P1 distance for WT (**F**) Bar plots showing the percentage of frames against C-term/C-term distance or C-term/P1 distance for L167F/F305L/F8L mutant. (**G**) Scatter/cluster plot of WT (orange) and the L167F/F305L/F8L (dark green) mutant C-term-C-term and C-term/P1 distances.

For example, Omicron/A206T that conferred strong resistance to drug treatment, was a frequently occurring mutation in our experiments and is distant from the catalytic site, prompting us to investigate the structure in more detail. Parental Omicron has a hydrophobic side chain alanine residue buried inside the protein at position 206, which is surrounded by the hydrophobic side chains of L253, P293, and V296 (**Fig. 5A**). The larger, polar side chain of the threonine mutant would be incompatible with the hydrophobic local environment, sterically clashing with the residues nearby, thereby requiring conformational changes to accommodate the side chain of threonine.

We also looked at L167F/F305L/F8L in more detail since it has both an N-terminal and C-terminal cleavage site mutation as well as the catalytic site L167F. In the different M^pro^ assays, mutations F305L and F8L alone did not confer resistance. In addition, the F305L mutant showed a gain-of-function phenotype (**Fig. 3A**) (*34*). Consistent with these findings, we observed a considerable decrease in stability for L167F (+61.9 kcal/mol), a similar stability for F8L (+1.09 kcal/mol) and an increase in protein stability for F305L (-5.7 kcal/mol), compared to the WT.

To investigate the effect of F305L in more detail, dimerization affinity prediction calculations were performed using Osprey, version 3.3 (*50*). This program uses rotamers of residue side chains to provide an estimation of binding affinity between two interaction partners, with higher Log_10_ K* scores indicating stronger and lower Log_10_ K* scores weaker binding. An increase in the binding affinity of +9.3 in the F305L mutant was calculated compared to the WT, suggesting an increase of M^pro^ dimerization affinity and formation of the mature dimer. F8L, conversely, did not influence dimerization affinity (**Fig. 5C**). F8 and F305 interact with each other via a pi-pi T-stack (**Fig. 5D and S11D**). In addition, F305 might be affected by electronic repulsion between the aromatic ring of F305 with the backbone oxygen atoms of I152 on protomer A and S123 on protomer B. Upon mutation to L305, the shorter leucine side chain is located further away from the backbone, potentially reducing this electronic repulsion between both protomers A and B (**Fig. 5D**).

Moreover, the contemporary presence of the mutations F8L and F305L likely causes an increase in C-terminus flexibility that could compete with the ligand for the binding site by interfering with its entry or acting as a “wiper” by occasionally leaping towards the active site (**Fig. 5E-G**).

The predicted loss of stability was less severe for the M^pro^ L167F/F305L/F8L (+2.2 kcal/mol) mutant. The Omicron A206T/F8L (+6.1 kcal/mol) also experienced a mild destabilization. For the mutant O/A206T/T198I we predicted an increased stability (-18.1 kcal/mol), indicating that the effects of this mutation might be mediated through a different mode of action.

Besides their impact on drug sensitivity, many of the mutations that decreased protease stability also increased the TM_50_ (**Fig. 3C**), potentially indicating impaired protease activity. For this reason, the stability of all the generated mutants was calculated, in addition to affinity changes of residues within the dimerization interface, for the apo dimer (without nirmatrelvir) (**Fig. S13A,C**).

Overall, we predicted a decrease of protein stability for most mutations similar to the nirmatrelvir results. This is not surprising, given that nirmatrelvir binds to the active M^pro^ conformation and the apo and nirmatrelvir-bound structures are therefore very similar (RMSD of 0.89 Å). The dimerization affinity was less affected than protease stability, with most mutants having a predicted Log_10_ K* score similar to the WT (**Fig. S13B**). Notable exceptions include C128Y, G2D, and G124D which were predicted to disrupt the dimer interaction instead (**Fig. S13B**). This finding was consistent with in vitro experiments, where we failed to plaque-purify VSV-C128Y-M^pro^.

To analyze specific multiple-catalytic site mutants in more detail, we performed Thermal Titration Molecular Dynamics (TTMD) (*52*) simulations. This approach allows the characterization in high detail of the binding mode and residence time of a ligand within the catalytic site of a given protein-ligand complex. It thereby enables the comparison between: 1. different ligands bound on the same protein (*52*); 2. different ligand conformations within the same binding site; 3. different mutants of the same protein in complex with the same ligand, as in the present work. The simulation consists of a series of classic molecular dynamics simulations performed at progressively increasing temperatures. Whether the ligand remains bound or not, is monitored through a scoring function based on protein-ligand interaction fingerprints (IFP*CS*) (*53*). In the end, the simulation returns the so-called MS coefficient, which is a measure for how long the protein-ligand complex is stably formed. This measure facilitates a qualitative comparison of the different protein-ligand complex stabilities, for example, nirmatrelvir bound to different mutant M^pro^ variants. A schematic representation of TTMD is shown in **Fig. 6A**.

**Fig. 6.**
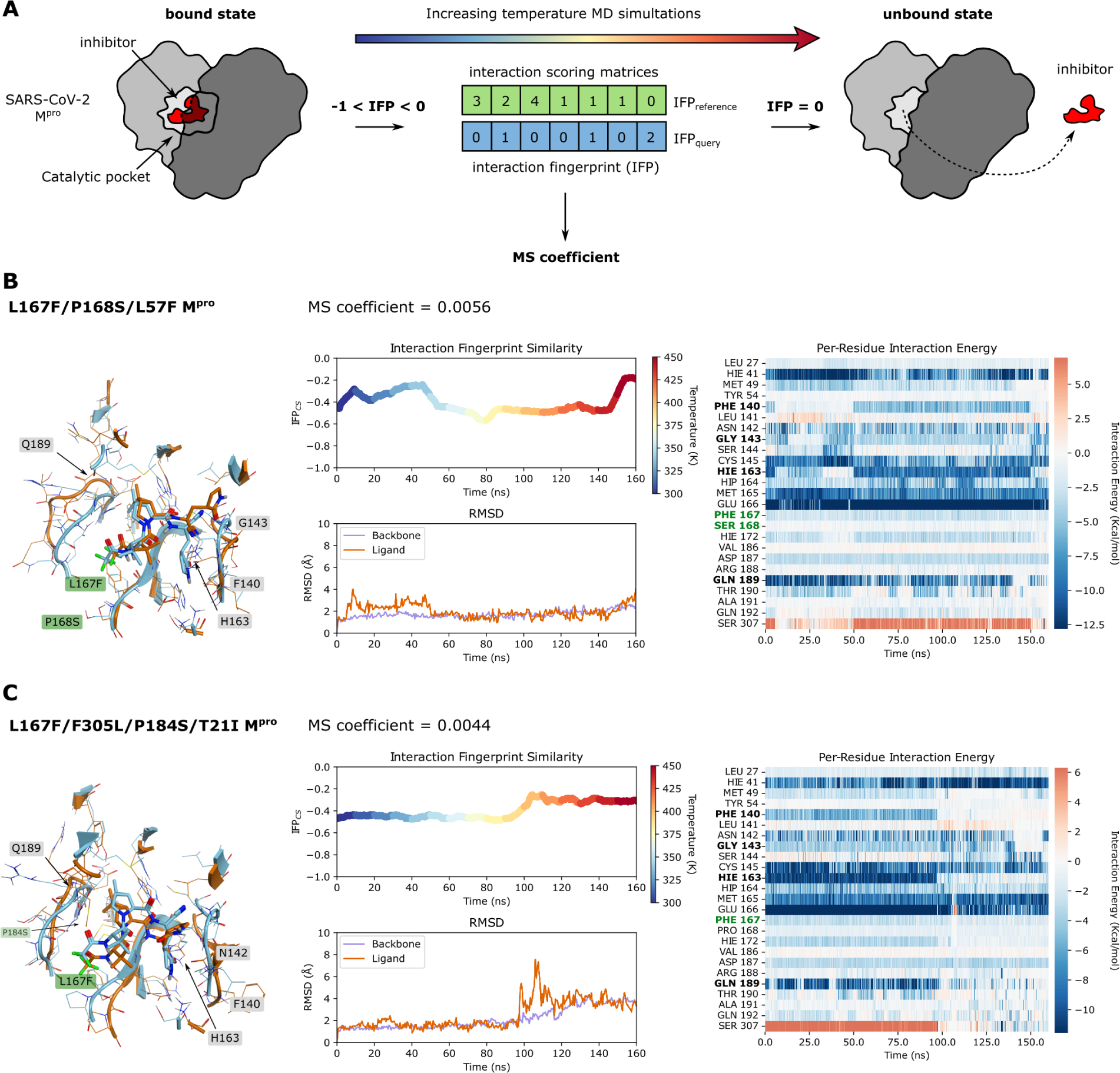
Thermal Titration Molecular Dynamics (TTMD) experiments of L167F/P168S/L57F and L167F/F305L/P184S/T21I. (**A**) Schematic representation of the TTMD simulation workflow. Interaction FingerPrints (IFP) of reference (bound state) and query (protease-inhibitor complex at each temperature condition over time) are continuously sampled and compared during the simulation. The closer from negative to zero the IFP_query_ is, the more the protease-inhibitor complex differs from the bound state. (**B**) TTMD simulation data of the nirmatrelvir-L167F/P168S/L57F- M^pro^ complex. Left: overlay of M^pro^ structure at the beginning (turquoise) and at the end of the simulation (orange); middle top: a rainbow plot including the IFP*CS* (left y-axis) is plotted against time in nanoseconds (x-axis). Additionally, the temperature in Kelvin is indicated by colours from blue to red on the right y-axis; middle bottom: the root-mean-square-deviation (left y-axis) is plotted against time in nanoseconds (x-axis); right: a heat map of interaction energies between ligand and surrounding residues. (**C**) TTMD simulation data of the nirmatrelvir- L167F/F305L/P184S/T21I-M^pro^ complex. Left: overlay of M^pro^ structure at the beginning (turquoise) and at the end of the simulation (orange); middle top: a rainbow plot including the IFP*CS* (left y-axis) is plotted against time in nanoseconds (x-axis). Additionally, the temperature in Kelvin is indicated by colours from blue to red on the right y-axis; middle bottom: the root-mean-square-deviation (left y-axis) is plotted against time in nanoseconds (x-axis); right: a heat map of interaction energies between ligand and surrounding residues. Residues that are mentioned in the results/discussion are highlighted in black, and mutated residues are highlighted in dark green.

Five catalytic site mutants (**Fig. 3A**) that arose from the WT-L167F-M^pro^ were selected for further investigation and compared to WT M^pro^-nirmatrelvir. Their structures, plus so-called rainbow blots, root-mean-square-deviations and interaction energy blots are depicted for the most resistant variants L167F/P168S/L57F and L167F/F305L/P184S/T21I (**Fig. 6B,C**).

Each TTMD simulation was repeated five times and the MS coefficients for each run are shown in **Table 1**. A plausible cutoff for the MS coefficient to discriminate between stable and unstable protein-ligand complexes is 0.004, according to recent literature (*52*, *54*). We observed that all the mutant MS values were above 0.004, meaning that the stability of the non-covalent nirmatrelvir- M^pro^ complex was lower for all of them compared to the WT protease. To illustrate the stabilities of the different complexes, we ranked the mutants according to MS coefficients, showing that the nirmatrelvir-WT complex is the most stable, followed by L167F/F305L/F8L > L167F/F305L/S144A > L167F/F305L/P184S/T21I > L167F/F305L/P184S and > L167F/P168S/L57F.

**Table 1.** MS coefficients of WT and mutants. MS coefficients were determined for each TTMD replicate, as well as the average MS value was calculated across three different replicates after discarding the highest and lowest values. The representative replicate (i.e., the closest to the average MS value), is highlighted with a star (*).

|  | Stability<br>rank<br>(decreasing) | Average | MD1 | MD2 | MD3 | MD4 | MD5 |
| --- | --- | --- | --- | --- | --- | --- | --- |
| Wildtype (WT) | 1 | 0.0037 | 0.0029 | 0.0028 | 0.0048 | 0.0037* | 0.0047 |
| L167F/P168S/L57F | 6 | 0.0056 | 0.0055* | 0.0069 | 0.0111 | 0.0038 | 0.0045 |
| L167F/F305L/S144A | 3 | 0.0044 | 0.0029 | 0.0044* | 0.0052 | 0.0042 | 0.0047 |
| L167F/F305L/P184S | 5 | 0.0051 | 0.0049 | 0.0071 | 0.0050* | 0.0054 | 0.0026 |
| L167F/F305L/F8L | 2 | 0.0042 | 0.0046 | 0.0041* | 0.0038 | 0.0039 | 0.0083 |
| L167F/F305L/P184S/T21I | 4 | 0.0045 | 0.0040 | 0.0044* | 0.0052 | 0.0049 | 0.0038 |

As can be observed in **Fig. 6B**, **Fig. S11B,C** and in **Movies S1-S3**, the active site constantly reshapes throughout the simulation, particularly in the area affected by the catalytic site mutations. L167F/P168S/L57F leads to a loss of interaction with the backbone of crucial catalytic site residues (164–166) by altering the floor of the binding site, causing a drag effect on the ligand (**Fig. 6B** and **Movie S1**). The resistance mechanism of the L167F/F305L/S144A mutant seemed to be increased instability of the oxyanion loop (residues 138 to 145), which finally leads to the loss of the hydrogen bond with H163 subsequently resulting in the loss of interaction with residue Q189 (*55*) (**Fig. S11B** and **Movie S2**). In L167F/F305L/P184S, the P184S mutation enhances the plasticity of the loop spanning from residue S184 to Q192, thereby reducing the ligand-pocket shape complementarity, which is essential for ligand positioning. At the same time, it did not alter the S1 pocket and the hydrogen bonds network within the central portion of the cavity (**Fig. S11C** and **Movie S3**). In mutant L167F/F305L/P184S/T21I the TTMD indicated a stabilizing effect of the T21I mutation (**Fig. 6C**, **Movie S5**). L167F/F305L/F8L (**Fig. S11D**) has only one mutation within the catalytic site (L167F) and two other ones located at the dimeric interface, F8L and F305L. As mentioned above, the C-terminus is freer to move in the available space, potentially competing with the ligand (**Movie S6**).

Further modeling results on the potential mechanisms leading to the resistance phenotype can be found in **Fig. 6B,C**; **Fig. S11A-D**, **Fig. S12,** and in **Movies S1-S6**.

## Discussion

In this study, we used a previously developed, safe, VSV-based tool to select for protease inhibitor resistance mutations. We demonstrated the capability of our selection tool to achieve SARS-CoV- 2 M^pro^ variants with multiple substitutions and potential Omicron-related mutants relevant to the current pandemic situation. The Omicron-M^pro^ sequence has a signature substitution at amino acid position 132, exchanging proline for histidine (P132H). This mutation has been shown to not confer resistance to approved protease inhibitors, such as nirmatrelvir and ensitrelvir (*29–32*), in agreement with our findings, but altered thermal stability (*33*). To reduce biosafety risks, previous studies based on authentic SARS-CoV-2 to select resistance mutations have used early variants of SARS-CoV-2 (*17–19*). Protease inhibitor resistant mutants derived from such early variants would unlikely be able to compete with Omicron variants if released accidentally. This caveat does not apply to mutation selection performed with our VSV-M^pro^ system. Consequently, we selected protease inhibitor resistant Omicron M^pro^ variants.

First, we generated a large number of mutants, which we subsequently filtered on a three-parameter basis for further characterization: proximity to the inhibitor, prevalence in the GISAID database, and frequency of a specific mutation in our selection experiments. Substitutions that appeared close to the inhibitor are likely to yield a meaningful, structural explanations that can then inform structural derivatization of inhibitors. Highly frequent substitutions in GISAID are relevant because they occur in viruses that can efficiently propagate in humans and therefore should not be detrimental to the enzymatic activity. Lastly, mutations that occur frequently in selection experiments could be favoured routes of evolution for the M^pro^ enzyme (*56*) and thus the mechanism might be of interest.

### Wuhan-1 M^pro^ resistance mutations

In the VSV-L167F-M^pro^ variant, additional substitutions close to the inhibitor were selected, namely S144A, P184S and L57F. In some of those mutants, we observed differences in the susceptibility to the two protease inhibitors nirmatrelvir and ensitrelvir. Interestingly, the mutant L167F/F305L/F8L demonstrated a considerably greater degree of resistance to ensitrelvir than to nirmatrelvir. Both L167F/F305L/P184S and L167F/F305L/L57F also showed a similar trend. L167F/P168S/S144A had a comparable resistance profile against both protease inhibitors. Mutations of S144 can simultaneously cause an increase in drug resistance and a decrease in the catalytic activity of the protease. Consistently, in our kinetic experiments and in the literature (*17*) this mutation impaired replication.

As mentioned above, the second filtering parameter we applied was the absolute frequency of a specific nsp5/M^pro^ substitution in the GISAID database. Residues in catalytic sites tend to be more conserved (*40*, *41*), and we expected them to not be frequently represented. Indeed, most high- frequent mutants were observed in positions further away from the active pocket. P184S is an exception as it is frequently represented (4713 entries, as of 01/06/2023) and is near the catalytic pocket. During the rescue (generation of a virus from a plasmid) of L167F/F305L/P184S, one sample had acquired an additional mutation, namely T21I. The resistance profile of L167F/F305L/P184S changed from more resistant against ensitrelvir to stronger resistance against nirmatrelvir upon introduction of the T21I mutation. However, T21I alone neither conferred resistance against nirmatrelvir nor ensitrelvir in both our gain- and loss-of-signal assays. Consistently, it has been reported in the literature as a compensatory mutation for the fitness loss caused by other resistance mutations, such as E166V (*17*). T21I could support or compensate the increased plasticity of L167F/F305L/P184S we found in TTMD simulations by an allosteric, re-stabilizing effect (*26*, *57*).

Taken together, it seems that L167F, P168S, S144A and P184S mutations deform the binding pocket and loosen the electrostatic interactions between nirmatrelvir and the protease. This observation indicates that not only mutations of residues directly interacting with the inhibitor can decrease susceptibility, but mutations of residues within the catalytic pocket can trigger conformational changes in the enzymatic cleft that impact in the inhibitor-protease complex formation as well.

The triple mutant L167F/F305L/F8L is among our most resistant M^pro^ variants and showed the most marked difference in susceptibility between the two inhibitors nirmatrelvir and ensitrelvir. The residues F305 and F8 are in close contact with each other in the folded protein and this is expected to be a stabilizing interaction for the dimer interface. However, other residues are viable at this position, for example leucine (L) has been shown to be the preferred residue instead of phenylalanine (F) (*40*). In fact, we calculated an increase in the dimerization affinity from F305 to L305, suggesting an improved formation of the fully functional dimeric protein upon mutation. We hypothesize that the simultaneous mutation of both F305 and F8 residues into leucines results in a modified interaction between the C-term of the first protomer and the N-term of the second, increasing the conformational degree of freedom of the C-term, thereby leading to a “wiper” mechanism that could compete with the inhibitor for the catalytic site (**Movie S6**).

### Omicron-M^pro^ resistance mutations

In the Omicron-M^pro^, we selected resistance mutations within and outside the catalytic site. The catalytic site mutant O/Y54H, which occurred 10 times independently, and O/R188W showed only mild resistance alone as well as in combination with R222L. O/Y54H/F305L showed an increased, selective resistance against ensitrelvir, with a 20- fold-change of the IC_50_ in the loss-of-signal assay (M^pro^-Off). The mutants K100N, T198I, A210S, and A266T were frequent in GISAID, therefore we picked them for resistance characterization. Double mutants O/A206T/F8L and O/A206T/T198I showed strong selective resistance against nirmatrelvir (up to 100-fold-change), while being less effective for ensitrelvir. We speculate that in O/A206T/F8L, the contribution of F8L to resistance is the same as in mutant L167F/F305L/F8L (“wiper” effect). O/A206T and O/A206T/K100N also showed considerable resistance (∼26-fold), and the computed destabilization correlated with resistance.

Most Omicron and WT substitutions, however, occurred in the amino acid sequence that constitutes domain III (residues 200-306) of M^pro^. We chose to characterize the A206T mutation due to the unusually high frequency in the selected VSV-O-M^pro^ pool of mutants. A206T only fits our third filtering parameter: frequent selection in our experiments, as it is far from the catalytic site and scarcely represented in the GISAID database. Likewise, some mutations may confer an advantage for viral replication only in the presence of a protease inhibitor, but would probably be unfavourable in an untreated individual. In a study on SARS-CoV M^pro^ polyprotein maturation (*58*), the authors proposed that despite the presence of deleterious mutations that hinder mature dimerization, M^pro^ retains its ability to cleave its N-terminal cleavage site owing to a weak, immature dimerization between two polyprotein monomers (1 and 2) catalysed by domain III (1) – domain III (2) interaction. We speculate that mutations of residues between 200 and 306, most frequently found in VSV-O-M^pro^, positively contribute to this interaction. As with A206T, we hypothesize that destabilization of the mature M^pro^ to which nirmatrelvir binds, is the resistance mechanism.

### Viral replication kinetics

In kinetic studies, we observed that the gain in signal was slower for some mutants, which might be an indirect read-out of M^pro^ activity. The L167F/F305L/S144A mutant was among the slowest variants and the time to reach the plateau was 1.75 times longer than for the WT M^pro^. However, mutations that were described as compensatory in the literature, such as T21I (*17*), could partially reverse the kinetic attenuation. T21I appeared after generating VSV-L167F-F305L-P184S-M^pro^ from its plasmid (also called “rescue”), and R222L after plaque purification of VSV-O-R188W-M^pro^. T21I and R222L are likely compensating for substitutions that occur in the catalytic site. For example, T21I seems to compensate for the loss of replicative fitness of the previously described, highly resistant M^pro^-E166V variant (*17*). Furthermore, T21I is a very frequently represented substitution in the GISAID database (21248 entries, 1^st^ of June 2023). From mutations generated in the VSV-O-M^pro^ variant, viral replication kinetics exhibited a different behaviour in comparison to VSV-L167F-M^pro^ variants. Overall, mutations related to the Omicron-M^pro^ did not decrease the viral replication rate to the same extent as in the L167F-M^pro^. The slowest variant was O/A206T/F8L, with a 1.4 fold-change (increase of ∼16 hours) to reach the plateau. As expected, most mutations that have been computationally predicted to decrease protease stability indeed had a negative impact on viral replication, with higher TM_50_ values related to M^pro^ variants bearing such mutations. Furthermore, they were less represented in isolates in the GISAID repository (1 or 2 digit entries), whereas those that had similar TM_50_ values compared to the WT such i.e. F8L, P184S or K100N and T198I alone, were more frequent (3 or 4 digit entries). These correlations support the validity of computational predictions.

### Caveats of the study

Chimeric VSV-Spike, where the VSV glycoprotein G was replaced by the SARS-CoV-2 spike, has been used previously to predict antibody escape mutations (*59–61*). Before that, VSV was shown many times to be promiscuous in the context of its glycoprotein, which could easily be replaced for example with those of Ebola and Marburg viruses (EBOV, MARV) (*62*) or lymphocytic choriomeningitis virus (LCMV) (*63*). To replace one of the VSV intergenic regions with a protease and thereby use VSV as a protease mutation tool, however, was a previously untapped area of research. Therefore, our resulting mutations need careful evaluation. We observed many mutations in chimeric VSV-M^pro^, some of which aligned exactly with those identified in SARS-CoV-2 gain-of-function experiments reported in the literature (*17*, *19*), whereas others were completely different. One explanation may be the artificial system used in our experiments, which uses cis-cleavage, the only requirement for M^pro^ processing in VSV-M^pro^ replication. In contrast, in experiments with authentic SARS-CoV-2, both cis- and trans-cleavage must be preserved in the development of resistance mutations. As described above, during auto- cleavage there is an intermediate dimerization state, different from the mature protease. This intermediate state may require specific interactions that differ from those involved in the mature dimer. Another reason is the difference between the natural polyprotein of coronaviruses and the artificial G-M^pro^-L polyprotein expressed by chimeric VSV-M^pro^. The observed divergence between SARS-CoV-2 and VSV may also be attributed to the differences between two viral polymerases: SARS-CoV-2 RNA-dependent RNA polymerase (RdRp) and VSV polymerase L. It has been reported in the literature that the SARS-CoV-2 polymerase can proofread (*64*), whereas VSV lacks such mechanisms (*36*), leading to a much higher error rate of 1/10,000 nucleotides, as described previously (*35*, *65*). However, this could also be seen as an advantage as resistance mutations may develop faster in the error prone VSV-M^pro^ replication system.

Overall, we conclude that most PI resistance mutations destabilize the mature conformation of M^pro^ and consequently the inhibitor-M^pro^ complex formation, which finally impairs inhibitor binding. These resistance mutations can be selective either for nirmatrelvir or for ensitrelvir, or impact both. Moreover, the extended use of protease inhibitors will increase the risk of selecting SARS-CoV-2 protease-inhibitor resistant variants. Therefore, to combat such variants, there is still a need for new PIs that ideally target alternative conformations or regions of the protease (*55*, *66*) and for rationale design of new inhibitors less affected by mutations (*67*).

## Materials and Methods

### Study Design

The overall rationale of the study was to use a previously developed mutation selection tool based on VSV to select and characterize a comprehensive collection of SARS-CoV-2 main protease inhibitor escape mutants. The study was performed on cell lines, in biochemical settings and in-silico, and no animal husbandry or human participants were involved. Human cell lines with replicating BSL-1 and -2 viruses were treated with protease inhibitors to observe resistance phenotypes in appropriate facilities. Viral titres were determined using TCID_50_. Measurement readouts were fluorescence-based, detected by ELISpot/FluoroSpot and multi-well readers. Autofluorescent fibers were excluded automatically and manually from spot counting in the ELISpot readout. Experiments were neither blinded nor randomly distributed to experimenters. We chose sample sizes empirically based on experience from former studies. At least two and up to four biologically independent replicates were performed per condition. Biologically independent meant distinct wells with the same condition, not multiple measurements of the same wells (technical replicates). Resistance phenotypes were reproduced at least twice, usually more often and in different combinations. Representative measurements were chosen to compile graphs and figures.

### Cloning strategies

The chimeric VSV variant with M^pro^ instead of the intergenic region between G and L was cloned as previously described (*34*). VSV-G-M^pro^-L carrying the Omicron signature mutation, P132H (VSV-O-M^pro^) was cloned as follows: N-terminal fragment comprising part of G and M^pro^ was amplified by PCR using primers 33n-before-KpnI-for (Q195) and Omicron rev VSV-G-M^pro^-L as a template, and the C-terminal fragment comprising the remaining part of M^pro^ and part of L was amplified by PCR using primers Omicron for and 33n-after-HpaI-rev (Q176). N- and C-terminal fragments were ligated together by Gibson assembly in a VSV-backbone vector digested with KpnI and HpaI. VSV-G-M^pro^-L L167F (VSV-L167F-M^pro^) was plaque purified as it was generated in a previous work.

M^pro^-Off and -On point mutants were generated by point directed mutagenesis on parental plasmids (GenBank accession codes: M^pro^-Off: ON262565; M^pro pro^-On: ON262564) with mutation primers and the Herculase II Fusion DNA Polymerase. Herculase is a polymerase that can overcome difficult (GC-rich) sequences and amplify large plasmids. However, primers could not be chosen for optimal design because the mutation site was fixed. For this reason, this simple point directed mutagenesis did not work for each construct. For M^pro^-On and M^pro^-Off plasmids, where point- directed mutagenesis did not work, we used mutagenic Gibson assembly as previously described (*34*). Cloning primers used in this study are shown in **Table S1**.

### Cell lines

BHK-21 cells (American Type Culture Collection, ATCC) were cultured in Glasgow Minimum Essential Medium (GMEM) (Lonza) supplemented with 10 % fetal calf serum (FCS), 5 % tryptose phosphate broth, and 100 units/ml penicillin and 0.1 mg/ml streptomycin (P/S) (Gibco). 293T cells (293tsA1609neo, ATCC), and 293-VSV (293 expressing N, P-GFP and L of VSV) (*68*) were cultured in Dulbecco’s Modified Eagle Medium (DMEM) supplemented with 10 % FCS, P/S, 2 % glutamine, 1x sodium pyruvate and 1x non-essential amino acids (Gibco). Vero E6 (ATCC CRL- 1586) was cultured in DMEM supplemented with 5 % FCS (VWR) and 1 % penicillin−streptomycin−glutamine (PSG) solution (Corning).

### Virus recovery (‘rescue’)

VSV-G-M^pro^-L WT and a few mutants were rescued in 293T cells by CaPO_4_ transfection of whole-genome VSV plasmids together with T7-polymerase, N-, P-, M-, G- and L expression plasmids as helper plasmids (*42*). Briefly, genome and helper plasmids were transfected into 293T in the presence of 10 µM chloroquine to avoid lysosomal DNA degradation. After 6 to 16 hours, chloroquine was removed, and cells were cultured until cytopathic effects occurred. M and G proteins were used as helper plasmids; although these proteins are optional in the recovery of VSV, they were chosen here as a precaution to support the rescue of a potentially attenuated virus variant. After the rescue, viruses were passaged on 293-VSV cells and plaque purified twice on BHK-21 cells. ΔP and ΔL VSV variants expressing dsRed were produced on replication supporting 293- VSV cells. VSV-G-M^pro^-L was fully replication competent and produced on BHK-21 cells.

### Mutation selection assay

10^4^ BHK-21 cells/well were seeded in a 96-well plate one day before infection with the chimeric VSV-M^pro^ viruses. VSV-L167F-M^pro^ and VSV-O-M^pro^ infected cells (MOI = 0.01) were treated with nirmatrelvir with concentration ranging from 10 to 100 µM. The resistant variants VSV- L167F/P168S-M^pro^ and VSV-L167F/F305L-M^pro^ at MOI = 0.01 were passaged 5 times, and with increasing concentrations of inhibitor (from 50 to 100 µM). VSV-O-M^pro^ and VSV-O/A206T-M^pro^ at MOI ranging from 0.001 and 0.01 were passaged only once with increasing concentrations of inhibitor (from 10 to 100 µM). Each virus variant occupied from 24 wells to 96 of the 96-well plate. Supernatants from wells that displayed cytopathic effect after 48-72 hours were collected for downstream viral RNA isolation, cDNA synthesis, PCR amplification and Sanger or Nanopore sequencing as described in this section.

### Viral RNA isolation, cDNA synthesis and M^pro^ sequencing

VSV-G-M^pro^-L RNA was isolated either by using the E.Z.N.A. Viral RNA Kit (Omega Bio-Tek Inc.), the NucleoSpin RNA Virus (Macherey-Nagel GmbH) or by semiautomated RNA extraction using the Easymag (BioMerieux^TM^). BHK-21 cells were infected with the respective VSV-G-M^pro^- L variant in 96-well plates as described above. Virus-containing supernatants were collected from individual 96-wells and the RNA was purified from the supernatants according to manufacturers’ instructions (Easymag). Then, cDNA was synthesized from isolated viral RNA by RevertAid RT Reverse Transcription Kit (Thermo Fisher Scientific). G_Cterm_-M^pro^-L_Nterm_ fragment sequence was amplified by PCR with primers (primer_for: CTCAGGTGTTCGAACATCCTCAC and primer_rev: GATGTTGGGATGGGATTGGC) and either sent for Sanger sequencing (MicroSynth AG) or sequenced using Nanopore (as described below). Obtained sequences were mapped to the L167F-M^pro^ or Omicron-M^pro^ reference sequences in Geneious Prime 2023.0.1 build 2022-11-28 and examined for mutations.

### Nanopore sequencing workflow

After the first two selection experiments where we sequenced our samples with Sanger sequencing (MicroSynth AG), we used Nanopore sequencing. First, PCR amplification products of the G_Cterm_- M^pro^-L_Nterm_ fragments, as described above, were used. The sequencing libraries were prepared using the Rapid Barcoding Kit SQK-RBK110.96 (Oxford Nanopore Technologies, ONT), and up to 96 samples were multiplexed and sequenced on a MinION Mk1B sequencer with R9.4.1 flowcells (ONT). Raw data in the form of electrical signals are translated into nucleotide sequences (base-calling) and saved in fastq files. Basecalling using the super high accuracy model, demultiplexing, as well as barcode and adapter sequence trimming, was performed in Guppy (version 6.1.5, ONT).

Raw reads were filtered to a PHRED quality score of ≥ Q15 and length between 200 bp and 1800 bp using SeqKit (version 2.4.0) (*69*), aligned to the reference sequence using minimap2 (version 2.22) (*70*), followed by sorting and indexing using SAMtools (version 1.13) (*71*). SAMtools depth was used to check sufficient depth. To find SNVs, we used LoFreq for variant calling with --min-cov option set to 300 (v2.1.3.1) (*72*).

The resulting VCF files were imported to Geneious Prime 2023.0.1 and called variants manually checked for plausibility.

### Unrooted phylogenetic tree generation and visualization

Mutations had to be manually inserted into the main protease sequence for each variant. The phylogenetic tree was generated using the online tool http://www.phylogeny.fr/ with standard parameters. The output as .newick file was imported to Geneious Prime 2023.0.1, visualized and exported as PDF. Graphic rearrangements and overhauls were made using Inkscape version 1.1.

### Collection of GISAID frequencies of generated mutations

To obtain updated nsp5 frequencies, metadata from the EpiCoV database (correct as of 1^st^ June 2023) was used; specifically, the ‘all_mutations’ field from a JSON dump obtained with GISAID credentials. Using Python 3.9 and the pyarrow and pandas’ libraries, sequences were classified into WT (not assigned Omicron lineage, and not carrying the P132H mutation) and Omicron (either assigned Omicron lineage, or carrying the P132H mutation). Then, for both these groups of sequences, the counts of 95 nsp5 mutations of interest were aggregated.

### SARS-CoV-2 variant phylogenetic analyses using Ultrafast Sample placement on Existing tRees (UShER)

Phylogenetic trees were generated for the resistance variants of interest using patient derived sequences deposited in the GISAID database. For each variant of interest, viral genomes harboring the corresponding mutation in Nsp5 were retrieved using the GISAID EpiCoV web server after filtering to consider only viruses descending from the Omicron lineage and removing genomes with low sequence coverage. These sequences were subsequently uploaded to the UShER (*73*) web server to generate phylogenetic trees using all the sequences available in the GISAID database, visualized using the Auspice.us web application and edited to highlight the variants of interest using Adobe Illustrator. The frequency of each resistance mutation within Omicron and Pre-Omicron lineages was similarly determined using the GISAID EpiCoV web server by dividing the number of occurrences of the corresponding Nsp5 mutation within Omicron or all other SARS-CoV-2 lineages by the total number of viral genomes deposited belonging to each respective lineage. Histograms comparing the frequency of the variants within the Omicron and Pre-Omicron lineages were generated using GraphPad Prism 9.

### Screening assay with ELISpot/FluoroSpot read-out

3 x 10^5^ cells were seeded per well in 6-well plates and transfected one day after seeding with 3CL^pro^ plasmids using TransIT-^PRO^ (Mirus Bio LLC) and incubated 8-9 hours for M^pro^-On and 10 hours for M^pro^-Off assays. Then, cells were seeded into a 96-well plate with 2 x 10^4^ cells per well in 50 µl complete growth medium. Directly after seeding, compounds and virus (MOI 0.1) were added in 50 µl complete growth medium to wells. After 48 to 72 hours, supernatants were removed, and fluorescent spots counted in a Fluoro/ImmunoSpot counter (CTL Europe GmbH). For experiments where FluoroBrite^TM^ medium was used, supernatants were not removed and the signal was read out every 12 hours. The manufacturer-provided software CTL switchboard 2.7.2. was used to scan 90% (70% when FluoroBrite^TM^ medium was used) of each well area concentrically to exclude reflection from the well edges, and counts were normalized to the full area. Automatic fiber exclusion was applied while scanning. The excitation wavelength for dsRed was 570 nm, and the D_F_R triple band filter was used to collect fluorescence. To increase comparability between M^pro^- On and -Off signals, we normalized dsRed events with the following strategies. In M^pro^-On, the highest compound concentrations would not reach the same value due to the different responses of each mutant. Therefore, we normalized to the highest mean of the experiment. In M^pro^-Off, we normalized the signal to each individual’s highest mean of the construct. For both assays, fluorescent spot count (y-axis) data were plotted against PI concentration (x-axis) and IC_50_ values were extrapolated as described later in this section.

### Replication kinetic experiments

For replication kinetic experiments with WT M^pro^, Omicron and mutants, we adapted the 3CL/M^pro^- Off assay to enable multiple read-outs over time every 12 hours. We exchanged the standard DMEM supplemented with 10 % FCS, P/S, 2 % glutamine, 1x sodium pyruvate and 1x non-essential amino acids (Gibco), with FluoroBrite^TM^ DMEM equally supplemented. FluoroBrite^TM^ DMEM has the same composition as DMEM, except it lacks phenol red which interferes with spot recognition. We experimented as described in the previous paragraph. After seeding 2 x 10^4^ transfected cells in 50 μl of FluoroBrite^TM^ complete medium into 96-well plates, only virus (MOI 0.1) was added (50 μl). The signal was read out at 12 hours intervals. As the virus starts replicating, dsRed is expressed by the infected cells, and the expression of dsRed is correlated with the amount of viral progeny and therefore with the number of spot count. We generally stopped signal acquisition after 84 up to 108 hpi, because the cells die due to virus-induced cytopathy, and the spot count decreases. The manufacturer-provided software CTL switchboard 2.7.2. was used as described above. Replication kinetic curves were normalized individually to the highest mean of the construct. We normalized fluorescent spot count data by the highest mean of each dataset. Then, we plotted the fluorescent dsRed signal (spot count) against time (hpi). We then performed non-linear regression analyses using the built-in function “Sigmoidal 4PL, X is concentration” in GraphPad Prism 9.5, where concentration was replaced with “hours” and extrapolated the TM_50_ (time-maximum fifty) value, which represents the time required to achieve half of the maximum signal/plateau.

### Protein Expression and purification

M^pro^ with a C-terminal Hexahistdine tag (M^pro^-6H) was expressed in *E.coli* followed by a tag removal with the human rhinovirus 14 3C protease (HRV3C protease).

WT M^pro^-6H construct was codon optimized for the expression in E. coli and ordered from Thermo Scientific (Waltham, MA, USA). Point mutations to generate the mutant variants were introduced using the Q5® Site-Directed Mutagenesis Kit purchased from New England Biolabs (NEB, Ipswich, MA, USA). Primers were acquired from Sigma Aldrich (St. Louis, MO, USA).

M^pro^-6H variants were transformed into electrocompetent *E. coli* BL21(DE3) cells. An overnight culture was inoculated 1:50 in 1L TB medium and cells were grown at 37°C, 220 rpm to an OD_600nm_ of 1. After induction with 0.4 mM Isopropyl β-D-1-thiogalactopyranoside (IPTG) temperature was lowered to 25°C for 4 hours. Cells were isolated by centrifugation at 3200 g, 20 minutes at 4°C, and cell pellets frozen at -20°C.

For purification, frozen pellets were resuspended in 50 mM Tris/50mM NaCl, pH 7.5 buffer and disrupted with a French press (Thermo Scientific). Lysed cells were centrifuged at 20000 g, 4°C, 20 min and the clarified supernatant was adjusted to 50 mM Tris/300 mM NaCl/20 mM Imidazole. IMAC purification was carried out with a ÄKTApurifier (GE Healthcare Life Sciences; Little Chalfon, United Kingdom) and a HisTrap FF Crude 1 mL (GE Healthcare) column. Unbound proteins washed out using 4 column volumes (CV) 50 mM Tris/300 mM NaCl/20 mM Imidazole, pH 7.5 and 1 CV 50 mM Tris/300 mM NaCl/40 mM Imidazole, pH 7.5. The protein-containing fractions were pooled, and the buffer was changed to 50 mM Tris/50mM NaCl, pH 7.5 using a HiTrap Desalting 5 mL (GE Healthcare) column. Protein concentration was determined with a NanoDrop ND-1000 Spectrometer (Thermo Scientific).

### Hexahistidine Tag removal by negative IMAC

The highly soluble Hexahistidine tagged NT*-HRV3Cprotease was expressed as previously described (*74*) and purified the same way as M^pro^. M^pro^-6H was incubated together with NT*- HRV3protease at 4°C overnight followed by negative IMAC purification with a manually packed Ni-NTA column. The flowthrough yields the mature M^pro^ with native N- and C-termini without His tag, because the His tag as well as the NT*-HRV3C protease are bound to the Ni-NTA resin.

### Cross validation with biochemical M^pro^ inhibition assay

The biochemical assay used to confirm mutations was based on the 3CL^pro^ activity assay from BPS Biosciences, catalog number #78042-2. The M^pro^ in the kit was replaced by an in-house produced M^pro^ and mutants thereof (L57F, L167F7P168S, L167F/P168S/L57F), as described in the M^pro^ purification paragraph. Solutions of WT and mutant M^pro^ variants – 84.5 ng/reaction in 30 µl buffer – (20 mM Tris/HCl pH = 8, 150 mM NaCl, bovine serum albumin 0.1 mg/ml, 1 mM dithiothreitol – DTT) were prepared according to the kit’s manual, to reach a final concentration of 50 nM in 50 µl. 10 µl of five-fold excess to tested nirmatrelvir concentrations (180 µM) were added to the 30 µl of M^pro^ solution and incubated for 30 minutes. Then, 10 µl of a fluorescent substrate (Ac-Abu-Tle-Leu-Gln-MCA) were added to get a final concentration of 40 µM. This generates a 1:5 dilution of the excess of protease inhibitors – and therefore final concentrations – and the reactions were incubated for 4 hours. Fluorescence was measured by excitation at 365 nm and read-out at 415- 445 nm emission with a Glomax Explorer fluorometer (Promega).

### Expression constructs

M^pro^ cutting reporter: Oligonucleotides for the “short” (GCAGTGCTCCAAAGCGGATTTCGC) and the “long” (ATCACGAGTGCAGTGCTCCAAAGCGGATTTCGCAAAATGGCC) cleavage sequence of M^pro^ (nsp4/nsp5 autocleavage site), flanked by N-terminal AgeI and a C-terminal HpaI restriction enzyme sites were ordered at Eurofins. The annealed oligos were ligated into a vector presenting N-terminally the fragment 1 (F[1]) and C-terminally the fragment 2 (F[2]) of the *Renilla* luciferase harboring the corresponding restriction enzyme sites (pcDNA3.1 backbone vector) as previously described (*75*, *76*). Between each fragment and the cleavage sequence we inserted interjacent 10-aa linkers.

### Preparation of lysates for western blotting

The cells expressing the M^pro^ cutting reporter and the purified proteins were prepared as described before. 50 µl of the protein suspension (5 µg of protein) were pre-incubated with inhibitor (10µM) or DMSO for 15 minutes at room temperature (R.T.). After that, 25 µl of cell suspension were added and the mixture was incubated for 3 hours at R.T. The cleavage reaction was stopped by the addition of 5x SDS-Loading buffer. Before the SDS-Page the samples were cooked at 95°C for 10 minutes. Western blot images were obtained using the ChemiDoc MP from Bio-Rad.

### TCID_50_ assay and dose responses

For initial dose response experiments, 5 x 10^4^ BHK-21 cells per well were seeded in 48-well plates one day before infection. Cells were infected in duplicates with a MOI of 0.05 of VSV-M^pro^ WT or VSV-O-M^pro^ and indicated concentrations of nirmatrelvir were added to the wells. After 48 hours, supernatants were collected and titrated with TCID_50_. For quantification, the 50 % tissue culture infective dose (TCID_50_) assay was performed as described previously (*77*). In short, 100 µl of serial dilutions of virus were added in octuplicates to 10^3^ BHK-21 cells seeded in a 96-well plate. Six days after infection, the TCID_50_ were read out and titers were calculated according to the Kaerber method (*78*).

### IC_50_ calculations

In this study, different assay systems were used to generate resistance data, namely VSV-based cellular assays with FluoroSpot read out, a luminescence-based cellular assay that employs recombinant M^pro^, as well as a peptide-MCA cleavage-based biochemical assay. IC_50_ calculations and statistical analysis for all assays were performed with GraphPad Prism 10. To calculate IC_50_s, we used GraphPads pre-set sigmoidal models: 4PL, X is concentration.

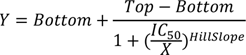

However, under some circumstances, slightly bell-shaped curves arise due to an initial signal in- or decrease (On and Off, respectively), followed by a signal peak and then a de- or increase. In the On assay, the decrease at the end occurs because of inhibitor toxicity at high concentrations. In the Off assay, the initial increase can be attributed to the fast death of cells in the absence of any inhibitor, whereas cells die slower at small doses, allowing for more cell division and thereby more “substrate” (cells and virus) generating dsRed. The bell-shaped curves can cause overfitting with the sigmoidal models, leading to extremely steep de- or increasing curves. Although in general, this does not lead to important changes in IC_50_s, we still chose to compensate for such unrealistically steep slopes by constraining the HillSlope to 3 or lower (positive slope) for the gain-of-signal assay and minus 3 or higher (negative slope) for the loss-of signal assay.

### Hardware overview (for subsequent TTMD simulations)

Most of the molecular modeling operations, such as the structure preparation, the setup for molecular dynamics (MD) simulations, and trajectory analyses, were performed on a Linux workstation, equipped with a 20 cores Intel Core i9-9820X 3.3 GHz processor, running the Ubuntu 20.04 operating system. Molecular dynamics simulations were carried out on an in-house Linux GPU cluster composed of 20 NVIDIA devices ranging from GTX1080Ti to RTX3090.

### Structure preparation

A computational study was conducted to rationalize the effect of SARS-CoV-2 M^pro^ mutations on Nirmatrelvir resistance. In detail, a recently developed enhanced sampling molecular dynamics method named Thermal Titration Molecular Dynamics (TTMD) (*52*, *54*) was exploited to compare the stability of the non-covalent complex between Nirmatrelvir and both the WT and mutated proteases, based on the assumption that the formation of the reversible, non-covalent complex, is the rate-determining step that leads to the formation of the irreversible (or very slowly reversible), covalent complex (*79*). The top five mutants concerning their experimentally determined drug resistance were investigated.

The experimentally determined three-dimensional coordinates of the WT nirmatrelvir-M^pro^ complex were retrieved from the Protein Data Bank (PDB) (*80*) and prepared for further calculations exploiting various tools provided within the Molecular Operating Environment (MOE) 2022.02 suite (*81*). Particularly, the complex deposited with accession code 7RFW 6 was used for describing the WT protease, while mutants were generated through the “protein builder” module of MOE.

The catalytically competent dimeric conformation of the protease was reconstructed by applying symmetric crystallographic transformations. Inconsistencies in the experimental structure were checked and fixed through the “structure preparation” tool. Missing hydrogen atoms were added according to the predominant tautomeric and protomeric state of each residue at pH = 7.4 through the “Protonate3D” module. Finally, each water molecule was removed before storing the prepared complex structure for subsequent calculations. In the end, a 1:1 protein-ligand non-covalent complex between nirmatrelvir and various M^pro^ proteases was considered in the simulated systems.

### System setup for MD simulations and Equilibration Protocol

Each complex obtained in the previous step was prepared for MD simulation making use of various tools provided within Visual Molecular Dynamics 1.9.2 (VMD) (*82*) and the AmberTools 22 (*83*, *84*) suite. All parameters were attributed according to the ff14SB (*85*) force field, except for the ligand ones which were instead assigned through the general Amber force field (GAFF) (*86*). Partial charges for the ligand were calculated through the AM1-BCC method (*87*). Each one of the protein-ligand systems was solvated using the TIP3P (*88*) water model into a rectangular base box, ensuring a minimum 15 Å distance between the box border and the nearest protein or ligand atom. To neutralize the net charge of the system and to reach a salt concentration of 0.154 M, the proper number of sodium and chlorine ions was added. Finally, to remove clashes and unfavorable contacts, 500 steps of energy minimization through the conjugate gradient algorithm were performed.

Before the production phase, a two-step equilibration process was performed. First, 0.1 ns of simulation in the canonical ensemble were carried out, imposing harmonic positional restraints on both the protein and ligand atoms. Then, 0.5 ns of simulation in the isothermal-isobaric ensemble were performed, with harmonic positional restraints applied only on the protein backbone and ligand atoms. In each case, a 5 kcal mol−1 Å−2 force constant was imposed on each restrained atom for the whole duration of the simulation, and the temperature was maintained at a constant value of 300 K through a Langevin thermostat (*89*). In the second equilibration stage, the pressure was kept constant at 1 atm through a Monte Carlo barostat (*90*).

All MD simulations were conducted exploiting the ACEMD16 3.5 engine, which is based upon the open-source Python library for biomolecular simulations OpenMM (*91*). A 2 fs integration timestep was used, and the M-SHAKE algorithm was used to constrain the length of bonds involving hydrogen atoms. The particle-mesh Ewald method was used to compute electrostatic interactions, using cubic spline interpolation and a 1 Å grid spacing. A 9.0 Å cutoff was applied to the computation of Lennard-Jones interactions.

### Thermal Titration Molecular Dynamics (TTMD) simulations

As thoroughly described in the work of Pavan et al. (*52*), Thermal Titration Molecular Dynamics (TTMD) is an enhanced sampling MD protocol originally developed for the qualitative estimation of protein-ligand unbinding kinetics.

The TTMD workflow relies on a series of short classic molecular dynamics simulations, defined as “TTMD-steps”, performed at progressively increasing temperatures. The temperature increase is used to augment the kinetic energy of the system, thus shortening the simulation time required to observe protein-ligand unbinding events compared to classic MD simulations. To monitor the progress of the simulation, the conservation of the native binding mode is evaluated through a protein-ligand interaction fingerprint (PLIF) based scoring function (*53*). The protocol described hereafter is implemented as a Python 3.10 code relying on the NumPy, MDAnalysis (*92*, *93*), Open Drug Discovery Toolkit (ODDT) (*94*) and Scikit-learn packages. The code is released under a permissive MIT license and available free of charge at github.com/molecularmodelingsection/TTMD.

In detail, the user must define a “temperature ramp”, i.e., the number, the temperature, and the length of each “TTMD-step”. Consistently with previous works on the target (*52*, *54*), in this case, the starting and end temperatures were set at 300K and 450K respectively, with a temperature increase between each “TTMD-step” of 10K and the duration of each simulation window set at 10 ns. The extension of the temperature ramp was determined based on the conservation of the native fold of the protein throughout the simulation, monitored through the backbone RMSD.

### Structure preparation (for subsequent analyses with Bioluminate and Osprey)

The PDB entries 7ALI (*51*), 8DZ2 (*48*), and 7TLL (*49*) were used to investigate the impact of mutations on apo WT, nirmatrelvir bound WT and nirmatrelvir bound Omicron dimers. All structures were prepared with the default settings of the Protein Preparation Wizard in Maestro version 2022-3 (*95*) except that selenomethionines were converted to methionines and no water molecules were deleted. The variant mutations Omicron, Omicron+A206T, L167F, L167F+P168S and L167F+F305L were introduced manually into the entry 7ALI (*51*) L167F, L167F+P168S and L167F+F305L were introduced manually into 8DZ2 (*48*) and Omicron+A206T was manually introduced in 7TLL (*49*). All mutations were introduced in Maestro version 2022-3 (*95*, *96*). The effects of additional mutations found in these most prevalent variants were investigated with the methods described below. Please note, only the structure 7ALI (*51*) includes residue 305 in both chains, 8DZ2 (*48*) only includes residue 305 in chain B, and 7TLL does not include residue 305 at all. Structure visualization and figure creation were performed in PyMol (*97*) version 2.5.0. Protein alignment was conducted in Maestro release 2022-3.

### Stability prediction

The stability of all the mutations was calculated using the default settings of the residue_scanning_backend.py module in Bioluminate (*44–47*) release 2022-3. Mutations were either introduced individually or in combination in the two protomers.

### Affinity prediction

For the affinity predictions in Osprey version 3.3 (*50*, *98–100*) water molecules were not considered. Osprey calculates so called Log10 K* scores (*99*) which provide an estimation of binding affinity. The required yaml and frcmod input files were created as described in detail in the Guerin et al. STAR Protocol (*101*). For the dimer affinity calculations, chain B was considered as the ligand. The stability threshold was disabled, epsilon was defined as 0.03 and WT and mutant side chain conformations were calculated as continuously flexible. Osprey is an open source software and is available for free at https://github.com/donaldlab/OSPREY3.

## Acknowledgements

We thank Florian Krammer, Bernhard Rupp and Sebastiaan Werten for useful discussions. The MMS lab is grateful to Chemical Computing Group, OpenEye, and Acellera for the scientific and technical partnership. MMS lab thankfully acknowledges the support of NVIDIA Corporation with the donation of the Titan V GPU used for this research.

## Author contributions

E.H. conceived the initial concept. F.C., E.H., A.D., H.S., B.S., J.F. and D.v.L. designed the experiments. E.H. and F.C. conceived the cloning strategies. E.H., F.C. generated recombinant viruses. E.H., F.C., S.R. and L.K. cloned plasmids for resistance experiments. E.H. and F.C. performed selection and resistance experiments with VSV-based methods and biochemical assays. A. Sauerwein, A.H. and T.R. performed viral RNA isolation and Nanopore library preparation. D. Bante elaborated Nanopore sequencing data. B.S. provided recombinant Mpro with point mutations expressed in E. coli. J.F. performed western blot analyses. S.A.M. provided phylogenetic analyses. J.H. provided substitution frequencies from the GISAID database. F.C. and E.H. wrote the manuscript and designed figures. A.D., M.P., D. Bassani, S. Menin and S. Moro performed structural modelling. H.S. and T.K. performed stability and dimerization affinity calculations. S. Moro and T.K. supervised in silico computational analyses. R.S. supervised B.S. recombinant Mpro production and E.S. supervised J.K. cross-validation assay. All authors read and approved the final manuscript.

## Data and materials availability

All data associated with this study are in the paper or supplementary materials. Further data can be obtained by contacting. <u>Materials can be obtained by contacting listed Email addresses</u> and after completion of a materials transfer agreement.

