## Supplementary Material for "A comprehensive study of SARS-CoV-2 main protease (M^pro^) inhibitor-resistant mutants selected in a VSV-based system"

Francesco Costacurta *et al.*

* Corresponding author.

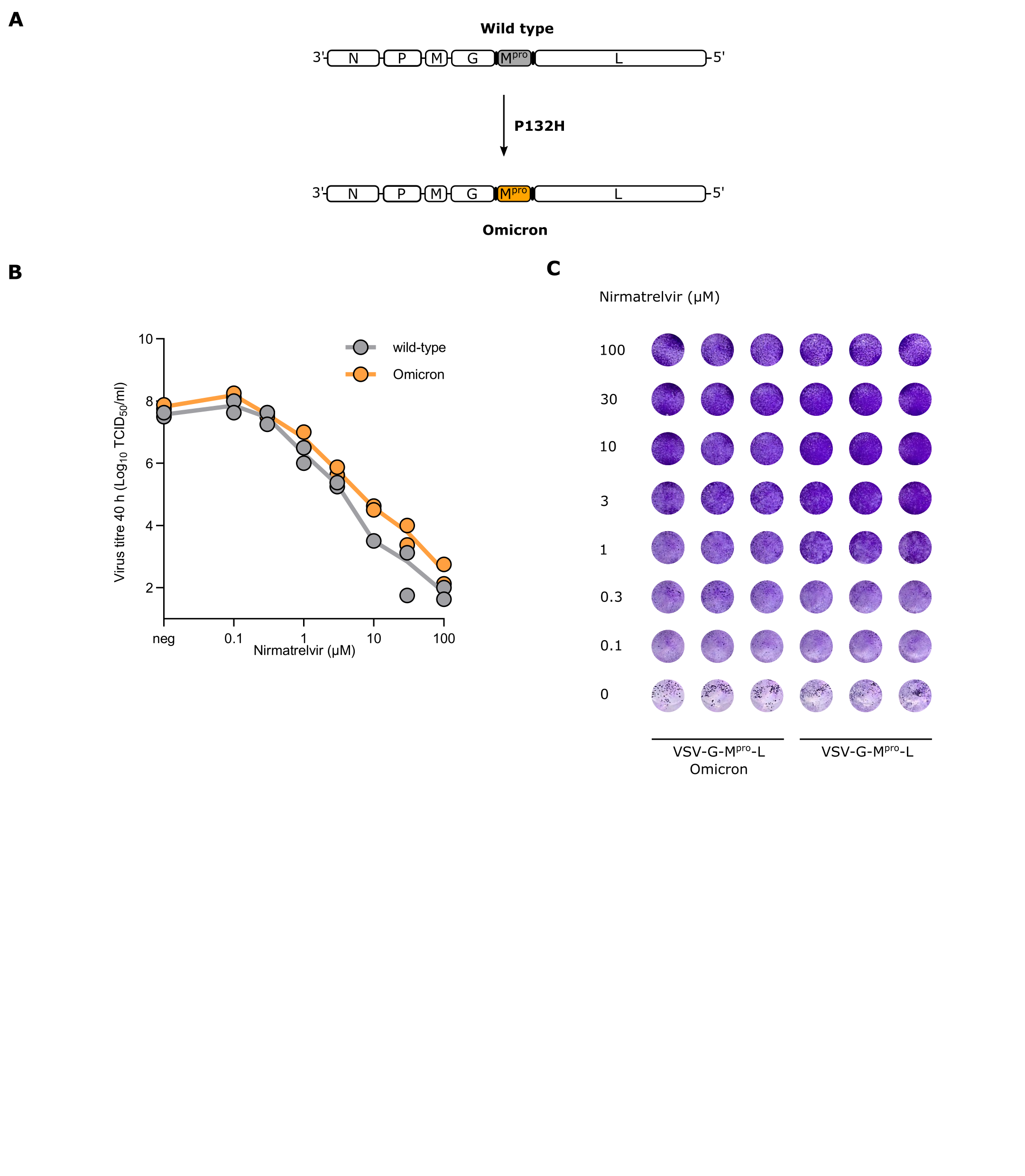

**Fig. S1. VSV-M^pro^ and VSV-Omicron-M^pro^ are equally susceptible to nirmatrelvir.** (**A**) Schematic representation of VSV-M^pro^ and VSV-Omicron-M^pro^ genomes. VSV-Omicron-M^pro^ was generated by introducing the substitution P to H at amino acid position 132 (P132H). (**B**) Dose response curves of VSV-M^pro^ and VSV-Omicron-M^pro^ against nirmatrelvir. Data are presented as the mean of n = 2 biologically independent replicates per condition. Each biological replicate consisted of n = 8 technical replicates. (**C**) Crystal violet staining of BHK21 cells used for dose response experiments with VSV-G-M^pro^-L. Data are presented as the mean of n = 3 biologically independent replicates per condition.

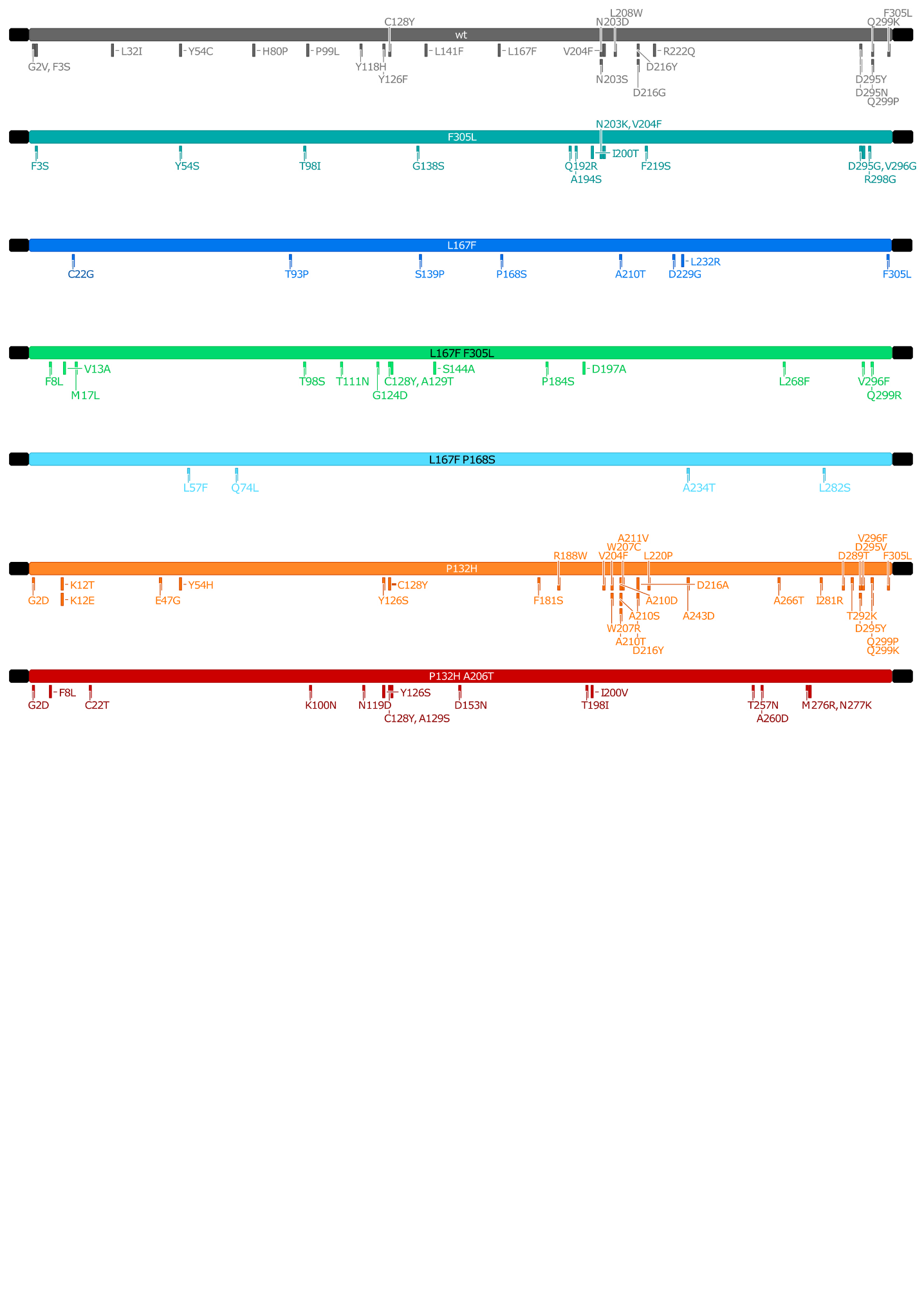
**Fig. S2. Different parental proteases and related mutations after selection experiments.** Schematic representation of the different parental proteases selected and used for further selection experiments with nirmatrelvir. From top to bottom: WT M^pro^ (Wuhan-1), F305L-M^pro^, L167F/P168S-M^pro^, L167F-M^pro^, L167F/F305L-M^pro^, P132H-M^pro^ (Omicron-M^pro^) and O/A206T-M^pro^. Mutations are represented by column-like symbols along the proteases, according to their position in the M^pro^ sequence.

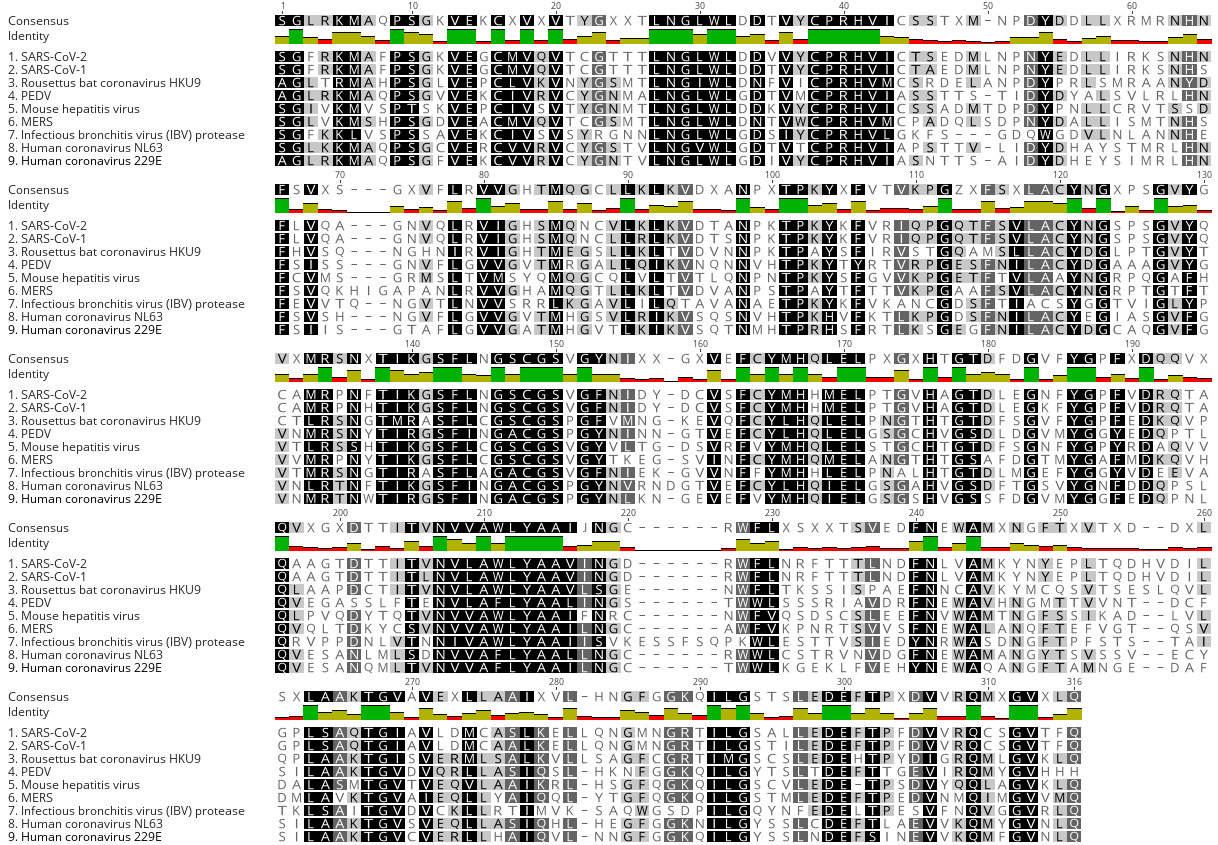

Fig. S3.

**Muscle alignment of M^pro^ and related coronavirus proteases.** MUltiple Sequence Comparison by Log-Expectation (MUSCLE) sequence alignment of SARS-CoV-2, SARS-CoV-1, Bat-CoV HKU9, PEDV, MHV (Mouse hepatitis virus), MERS, IBV, NL63 and 229E shows areas of conservation and amino acid variability.

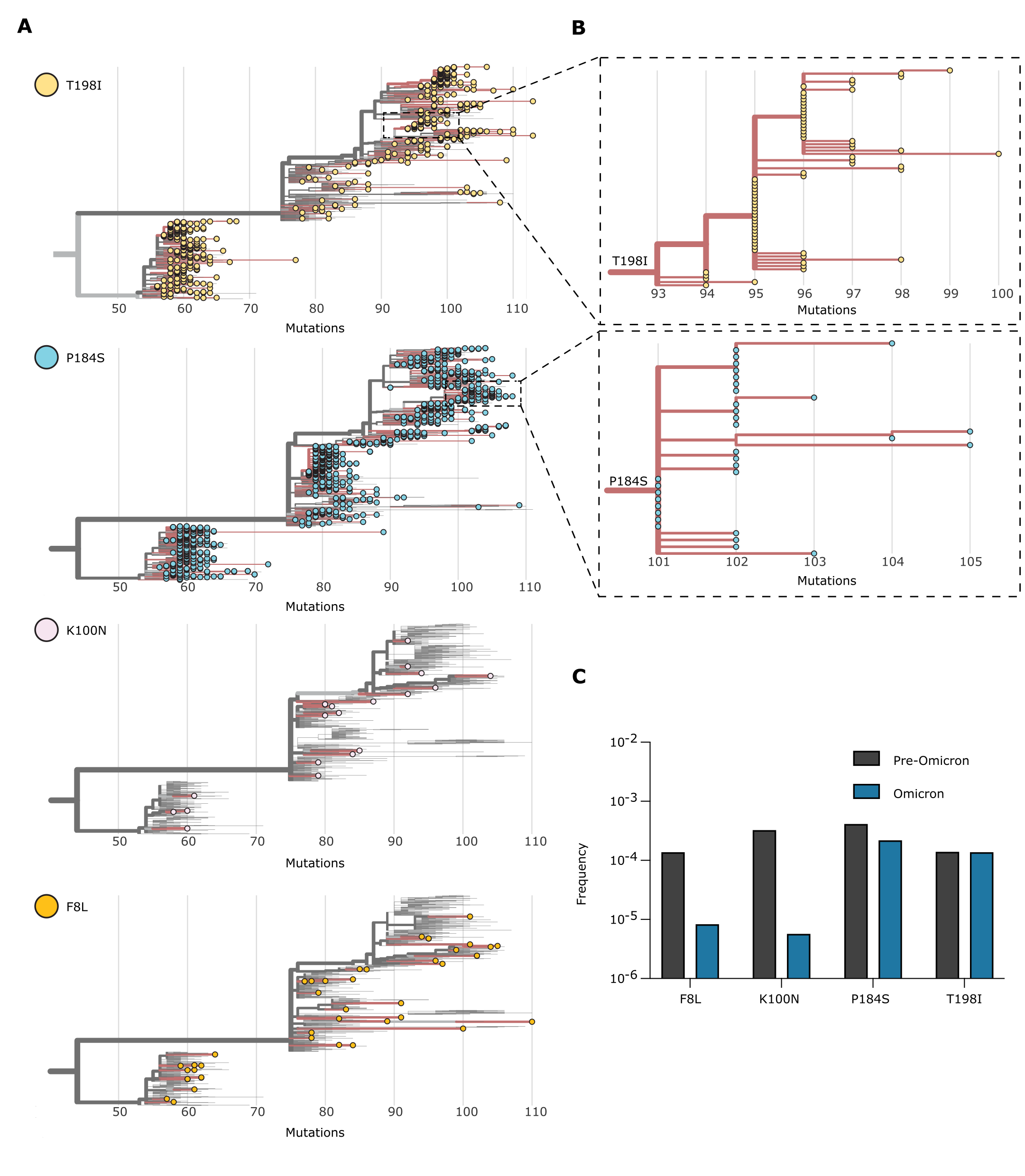

**Fig. S4. Phylogenetic subtrees of T198I, P184S, K100N, and F8L substitutions.** (**A**) Phylogenetic subtree of nsp5/M^pro^-T198I, -P184S, -K100N, -F8L substitutions generated with the Ultrafast Sample placement on Existing tRee (UShER) tool (GISAID, 18th January 2023). Only sequences deposited after the Omicron emergence were used. (**B**) Magnified view of the T198I and P184S subtree areas, showing transmission of this variant from a single founder event. (**C**) Frequency ratio of T198I, P184S, K100N, and F8L mutations compared with pre- and post-Omicron variant surge (GISAID, 18th January 2023).

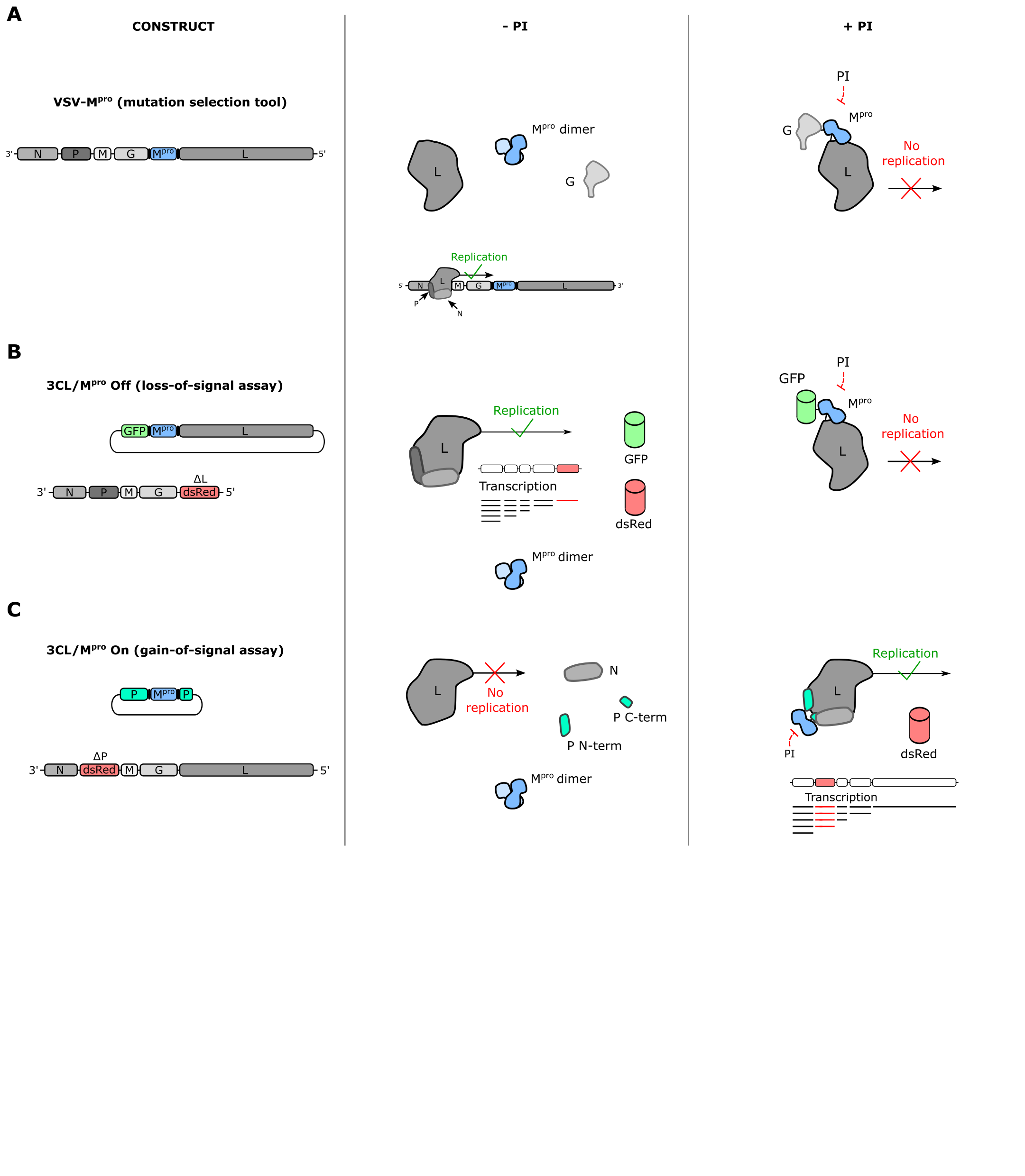

**Fig. S5. Comparison of the cell-based systems used in this study.** From left to right, schematic representation of: construct(s), mechanism in the absence of inhibitor, mechanism in the presence of inhibitor. (**A**) VSV-M^pro^ (mutation selection tool) construct and mechanism. Without inhibitor, M^pro^ processes G-M^pro^-L and the virus replicates. After an inhibitor is applied, M^pro^ is inactive and viral replication is blocked. (**B**) Graphical representation of the loss-of-signal cellular assay mechanism: VSV-ΔL-dsRed + G-M^pro^-L are added to cells. Without inhibitor applied, M^pro^ processes GFP-M^pro^-L and VSV-ΔL-dsRed replicates, expressing dsRed. After an inhibitor is applied, M^pro^ is inactive and VSV-ΔL-dsRed replication is turned off. (**C**) Graphical representation of the gain-of-signal cellular assay mechanism: VSV-ΔP-dsRed + P:M^pro^:P are added to cells. Without inhibitor, M^pro^ shatters the phosphoprotein (P) in two pieces, impairing viral replication of replication-incompetent VSV-ΔP-dsRed. After an inhibitor is applied, M^pro^ is inactive, P is intact and VSV-ΔP-dsRed replication is turned on and expresses dsRed.

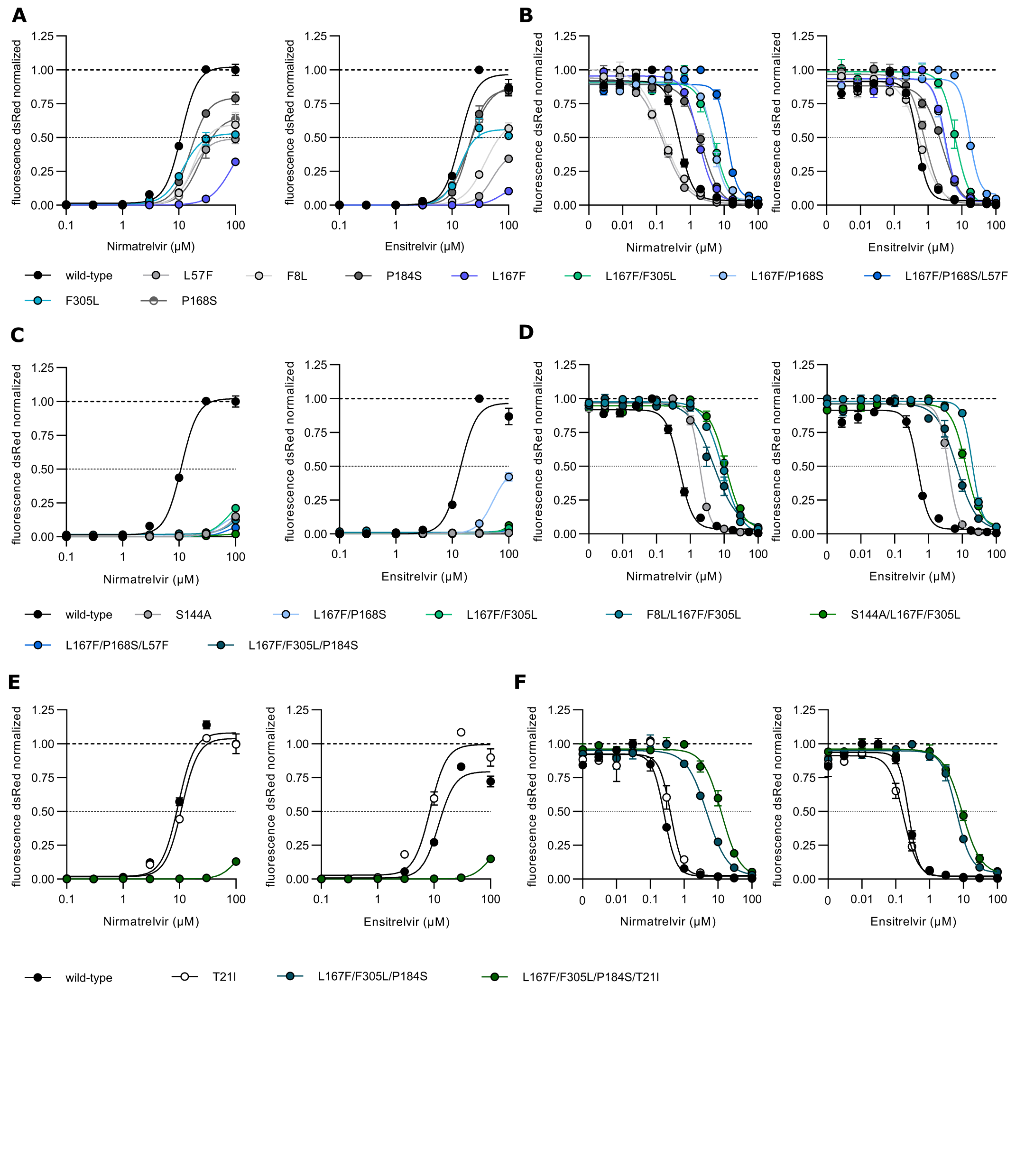

**Fig. S6. Dose response curve of WT and mutant main proteases.** (**A**) Gain-of-signal assay results are shown for M^pro^ WT and L167F, F305L, F8L, L57F, P184S, P168S mutants against the protease inhibitors nirmatrelvir (left) and ensitrelvir (right). Data are presented as SEM of n = 2 biologically independent replicates per condition. (**B**) Loss-of-signal assay results are shown for M^pro^ WT and F8L, L57F, L167F, P184S, L167F/P168S, L167F/F305L, L167FF/P168S/L57F mutants against the protease inhibitors nirmatrelvir (left) and ensitrelvir (right). The signal was read out at 48 hours post infection. Data are presented as SEM of n = 3/n = 4 biologically independent replicates per condition. Fold-change was calculated using the IC_50_ value of WT M^pro^ at this timepoint. (**C**) Gain-of-signal assay results are shown for M^pro^ WT and S144A, L167F/F305L, L167F/P168S, L167F/F305L/S144A, L167F/F305L/P184S, L167F/F305L/F8L, L167F/F305L/L57F mutants against the protease inhibitors nirmatrelvir (left) and ensitrelvir (right). Data are presented as SEM of n = 2 biologically independent replicates per condition. (**D**) Loss-of-signal assay results are shown for M^pro^ WT and S144A, L167F/F305L/F8L, L167F/F305L/S144A, L167F/F305L/P184S mutants against the protease inhibitors nirmatrelvir (left) and ensitrelvir (right). The signal was read out at 84 hours post infection. Data are presented as SEM of n = 3/n = 4 biologically independent replicates per condition. Fold-change was calculated using the IC_50_ value of WT M^pro^ at this timepoint. (**E**) Gain-of-signal assay results are shown for M^pro^ WT and T21I, L167F/F305L/P184S/T21I mutants against the protease inhibitors nirmatrelvir (left) and ensitrelvir (right). Data are presented as SEM of n = 2/n = 3 biologically independent replicates per condition. (**F**) Loss-of-signal assay results are shown for M^pro^ WT and T21I, L167F/F305L/P184S, L167F/F305L/P184S/T21I mutants against the protease inhibitors nirmatrelvir (left) and ensitrelvir (right). The signal was read out at 72 hours post infection. Data are presented as SEM of n = 3/n = 4 biologically independent replicates per condition. Fold-change was calculated using the IC_50_ value of WT M^pro^ at this timepoint.

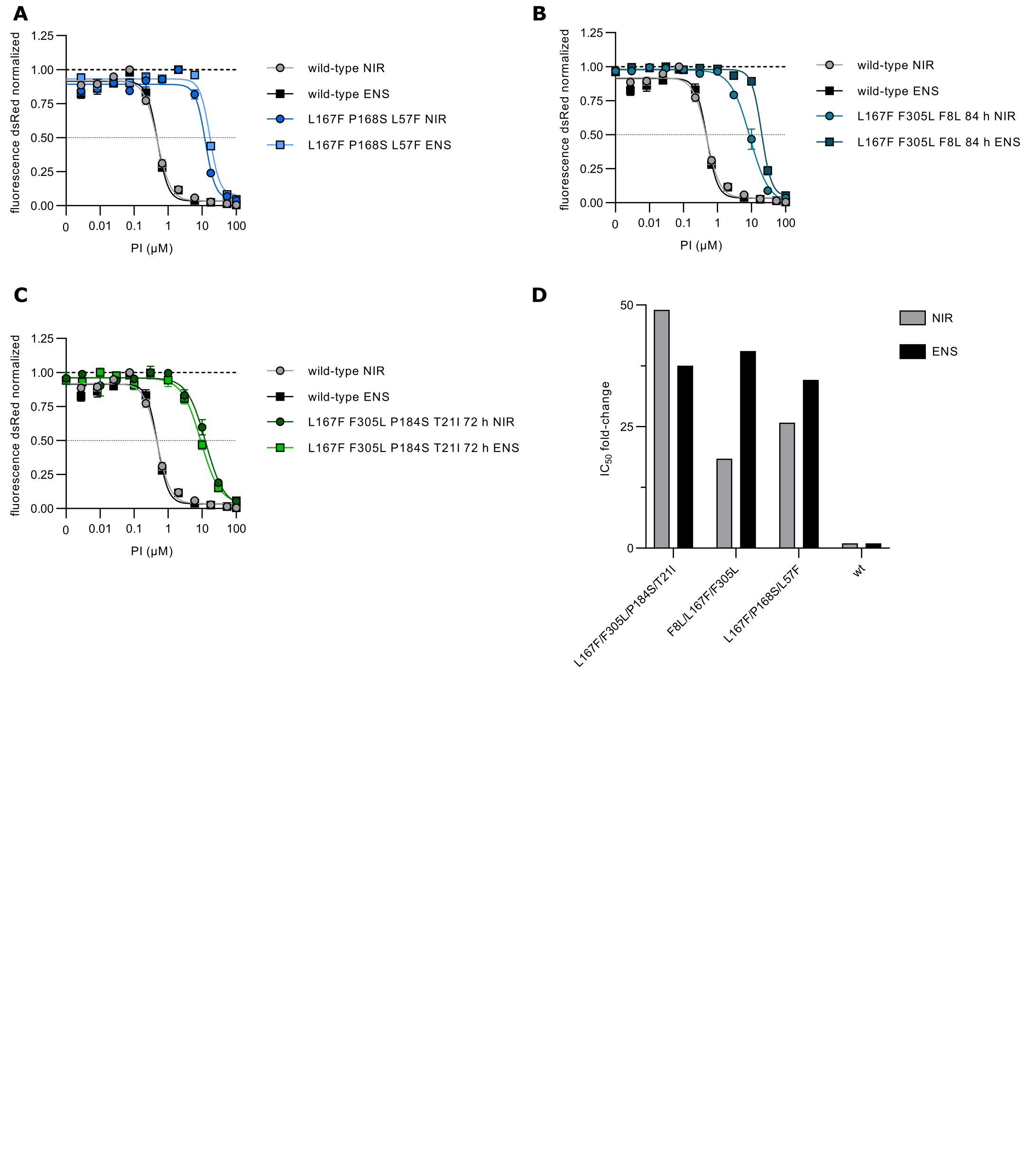

**Fig. S7. IC values shifts in M^pro^-Off WT** **mutants.** (**A**) Loss-of-signal assay results are shown for WT M^pro^ and L167F/P168S/L57F mutant against the protease inhibitors nirmatrelvir and ensitrelvir. The signal was read out at 48 hours post infection. Data are presented as SEM of n = 3/n = 4 biologically independent replicates per condition. (**B**) Loss-of-signal assay results are shown for WT M^pro^ and L167F/F305L/F8L mutant against the protease inhibitors nirmatrelvir and ensitrelvir. The signal was read out at 48 hours post infection. Data are presented as SEM of n = 3/n = 4 biologically independent replicates per condition. (**C**) Loss-of-signal assay results are shown for WT M^pro^ and L167F/F305L/P184S/T21I mutant against the protease inhibitors nirmatrelvir and ensitrelvir. The signal was read out at 48 hours post infection. Data are presented as SEM of n = 3/n = 4 biologically independent replicates per condition. (**D**) Barplot representing the IC50 fold-changes of L167F/P168S/L57F, L167F/F305L/F8L and L167F/F305L/P184S/T21I mutants compared to the WT for nirmatrelvir and ensitrelvir.

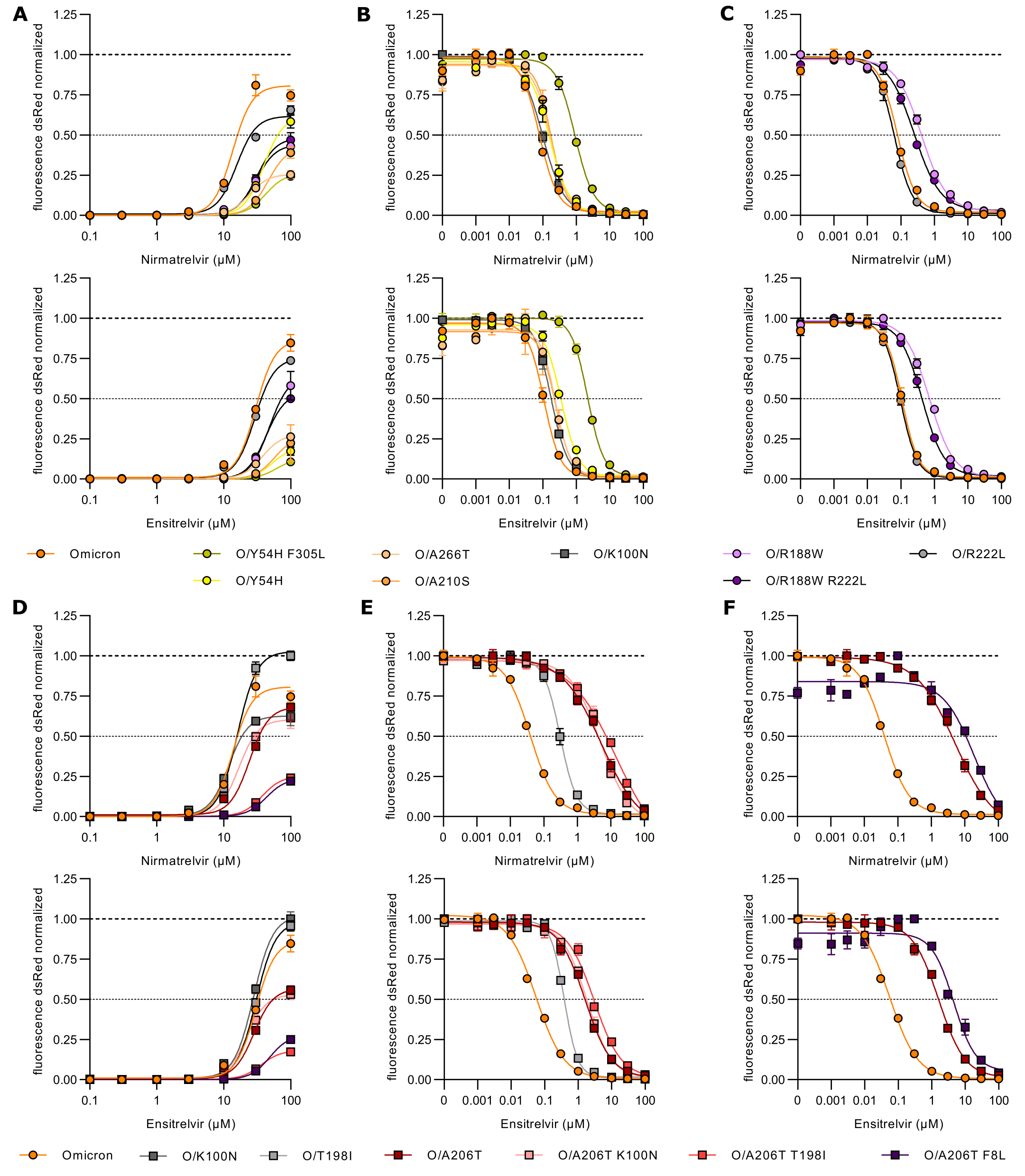

**Fig. S8. Dose response curve of Omicron and mutant main proteases.** (**A**) Gain-of-signal assay results are shown for Omicron-M^pro^ and O/A266T, O/A210S, O/Y54H, O/Y54H+F305L, O/R188W, O/R222L, O/R188W/R222L mutants against the protease inhibitor nirmatrelvir (top) and ensitrelvir (bottom). Data are presented as SEM of n = 3 biologically independent replicates per condition. (**B**) Loss-of-signal assay results are shown for Omicron-M^pro^ and O/A266T, O/A210S, O/Y54H, O/K100N, O/Y54H+F305L mutants against the protease inhibitor nirmatrelvir (top) and ensitrelvir (bottom). The signal was read out at 48 hours post infection. Data are presented as SEM of n = 4 biologically independent replicates per condition. Fold-change was calculated using the IC_50_ value of Omicron-M^pro^ at this timepoint. (**C**) Loss-of-signal assay results are shown for Omicron-M^pro^ and O/R188W, O/R222L, O/R188W/R222L mutants against the protease inhibitor nirmatrelvir (top) and ensitrelvir (bottom). The signal was read out at 48 hours post infection. Data are presented as SEM of n = 4 biologically independent replicates per condition. Fold-change was calculated using the IC_50_ value of Omicron-M^pro^ at this timepoint. (**D**) Gain-of-signal assay results are shown for Omicron-M^pro^ and O/A206T, O/T198I, O/K100N, O/A206T/F8L, O/A206T/K100N mutants against the protease inhibitor nirmatrelvir (top) and ensitrelvir (bottom). Data are presented as SEM of n = 3 biologically independent replicates per condition. (**E**) Loss-of-signal assay results are shown for Omicron-M^pro^ and O/A206T, O/T198I, O/A206T/T198I, O/A206T/K100N mutants against the protease inhibitor nirmatrelvir (top) and ensitrelvir (bottom). The signal was read out at 60 hours post infection. Data are presented as SEM of n = 3/n = 4 biologically independent replicates per condition. Fold-change was calculated using the IC_50_ value of Omicron-M^pro^ at this timepoint. (**F**) Loss-of-signal assay results are shown for Omicron-M^pro^ and O/A206T, O/A206T/F8L mutants against the protease inhibitor nirmatrelvir (top) and ensitrelvir (bottom). The signal was read out at 60 hours post infection. Data are presented as SEM of n = 3/n = 4 biologically independent replicates per condition. Fold-change was calculated using the IC_50_ value of Omicron-M^pro^ at this timepoint.

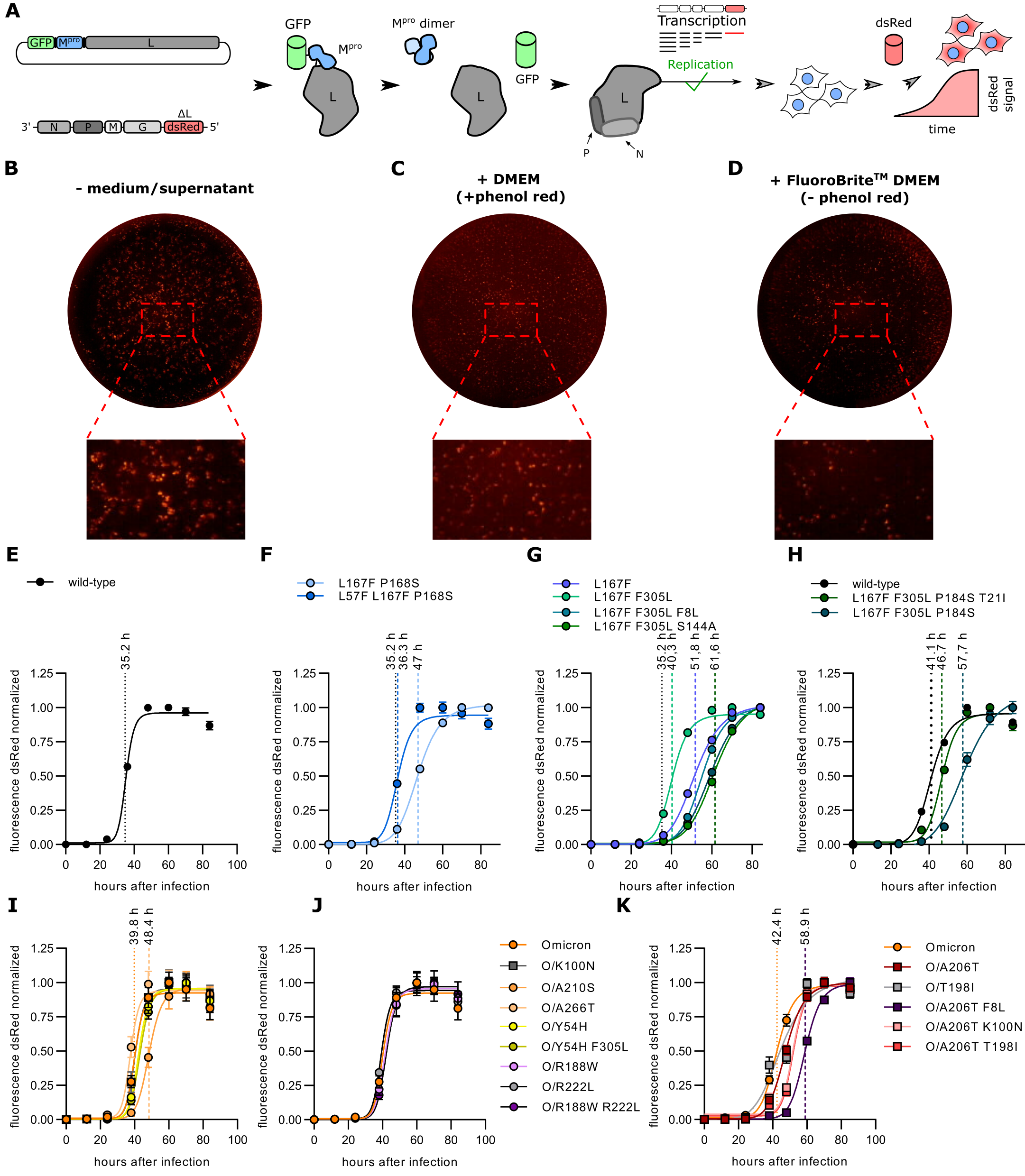

**Fig. S9. 3CL/M^pro^-Off kinetics mechanism & normal DMEM/FluoroBrite^TM^ DMEM culture media differences.** (**A**) Schematic representation of the M^pro^-Off assay adaptation to a viral replication kinetic measurement based on M^pro^ activity. M^pro^-Off transfected cells are infected with VSV-ΔL-dsRed and the signal is read out over time. No protease inhibitor is applied. (**B**) Representative photo, and magnified view (below), taken with the ELISpot reader (FluoroSpot X suite) of a well after removing the supernatant. (**C**) Representative photo, and magnified view (below), taken with the ELISpot reader (FluoroSpot X suite) of a well where the supernatant was not removed (DMEM, + phenol red). (**D**) Representative photo, and magnified view (below), taken with the ELISpot reader (FluoroSpot X suite) of a well where the supernatant was not removed (FluoroBrite^TM^ DMEM, - phenol red). (**E**) Replication kinetics fitting curve of WT M^pro^ (± SEM; n = 8 biological replicates). The dotted line represents the TM_50_ value related to the WT M^pro^. (**F**) Replication kinetics fitting curves of L167F/P168S and L167F/P168S/L57F M^pro^ mutants (± SEM; n = 8 biological replicates). The dotted lines represent the TM_50_ value related to the WT M^pro^ and the triple mutant L167F/P168S/L57F. (**G**) Replication kinetics fitting curves of L167F, L167F/F305L, L167F/F305L/P184S, L167F/F305L/F8L and L167F/F305L/S144A M^pro^ mutants (± SEM; n = 8 biological replicates). The dotted lines represent the TM_50_ value related to the WT M^pro^ and the mutants L167F, L167F/F305L, L167F/F305L/S144A. (**H**) Replication kinetics fitting curves of T21I, L167F/F305L/P184S, L167F/F305L/P184S/T21I M^pro^ mutants (± SEM; n = 8 biological replicates). The dotted lines represent the TM_50_ value related to the mutants T21I, L167F/F305L/P184S, L167F/F305L/P184S/T21I. (**I**) Replication kinetics fitting curves of Omicron, O/Y54H, O/Y54H/F305L, O/A210S, O/A266T, O/K100N M^pro^ mutants (± SEM; n = 8 biological replicates). The dotted lines represent the TM_50_ value related to the Omicron main protease and the mutant O/A210S. (**J**) Replication kinetics fitting curve of Omicron, O/R188W, O/R222L, O/R188W/R222L M^pro^ mutants (± SEM; n = 8 biological replicates). (**K**) Replication kinetics fitting curves of Omicron, O/A206T, O/A206T/F8L, O/A206T/T198I, O/A206T/K100N, O/T198I M^pro^ mutants (± SEM; n = 8 biological replicates). The dotted lines represent the TM_50_ value related to the Omicron main protease and the mutant O/A206T/F8L.

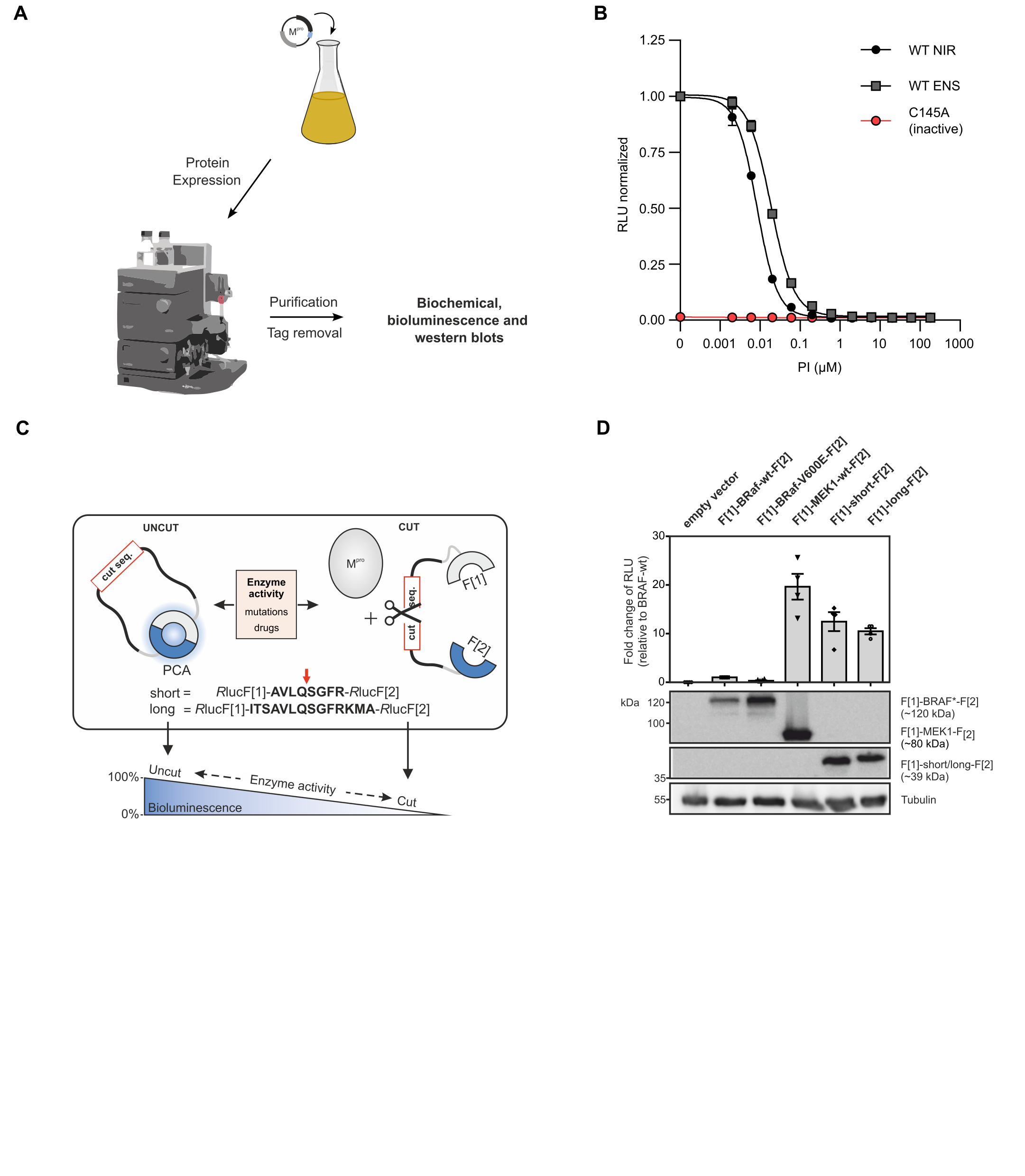

**Fig. S10. Selected recombinant proteases.** (**A**) Following recombinant protein expression, M^pro^ variants were purified using a FPLC system for subsequent assays employing the recombinant protease. (**B**) Dose response experiment of WT and C145A mutant (catalytically inactive) with nirmatrelvir and ensitrelvir (± SEM; n = 2). (**C**) Depiction of the PCA-based M^pro^-cutting reporter. The cutting sequence of M^pro^ is flanked by two fragments of the Renilla luciferase indicated with F[1] and F[2]. The intact M^pro^ cutting reporter allows both fragments to form a functional luciferase protein capable to emit light after substrate addition. Incubation of the reporter with purified M^pro^ leads to reporter cleavage and a subsequent signal decrease. Protease inhibitor binding and mutations can influence the cleavage event, thereby altering the PCA-emitted bioluminescence signals. (**D**) Bioluminescence signal strength test of the two M^pro^-cutting reporters (short; long). HEK293T cells expressing indicated bioluminescence reporter or empty vector were subjected to a bioluminescence measurement. Bars represent the fold change of emitted bioluminescence signals relative to the BRaf-wt reporter (±SEM; n = 3). Below a representative western blot is shown, revealing good and equal reporter expression between both generated cutting reporters.

**Fig. S11.** **Thermal Titration Molecular Dynamics (TTMD) experiments of WT, L167F/F305L/S144A, L167F/F305L/P184S and L167F/F305L/F8L.** TTMD simulation data of M^pro^ mutants. Left: overlay of M^pro^ structure at the beginning (turquoise) and at the end of the simulation (orange); middle top: a rainbow plot including the IFP*_CS_* (left y-axis) is plotted against time in nanoseconds (x-axis). Additionally, the temperature in Kelvin is indicated by colours from blue to red on the right y-axis; middle bottom: the root-mean-square-deviation (left y-axis) is plotted against time in nanoseconds (x-axis); right: a heat map of interaction energies between ligand and surrounding residues. Residues that are mentioned in the results/discussion are highlighted in black, and mutated residues are highlighted in dark green. (**A**) Wildtype (WT) M^pro^ (**B**) L167F/F305L/S144A mutant (**C**) L167F/F305L/P184S mutant (**D**) L167F/F305L/F8L mutant.

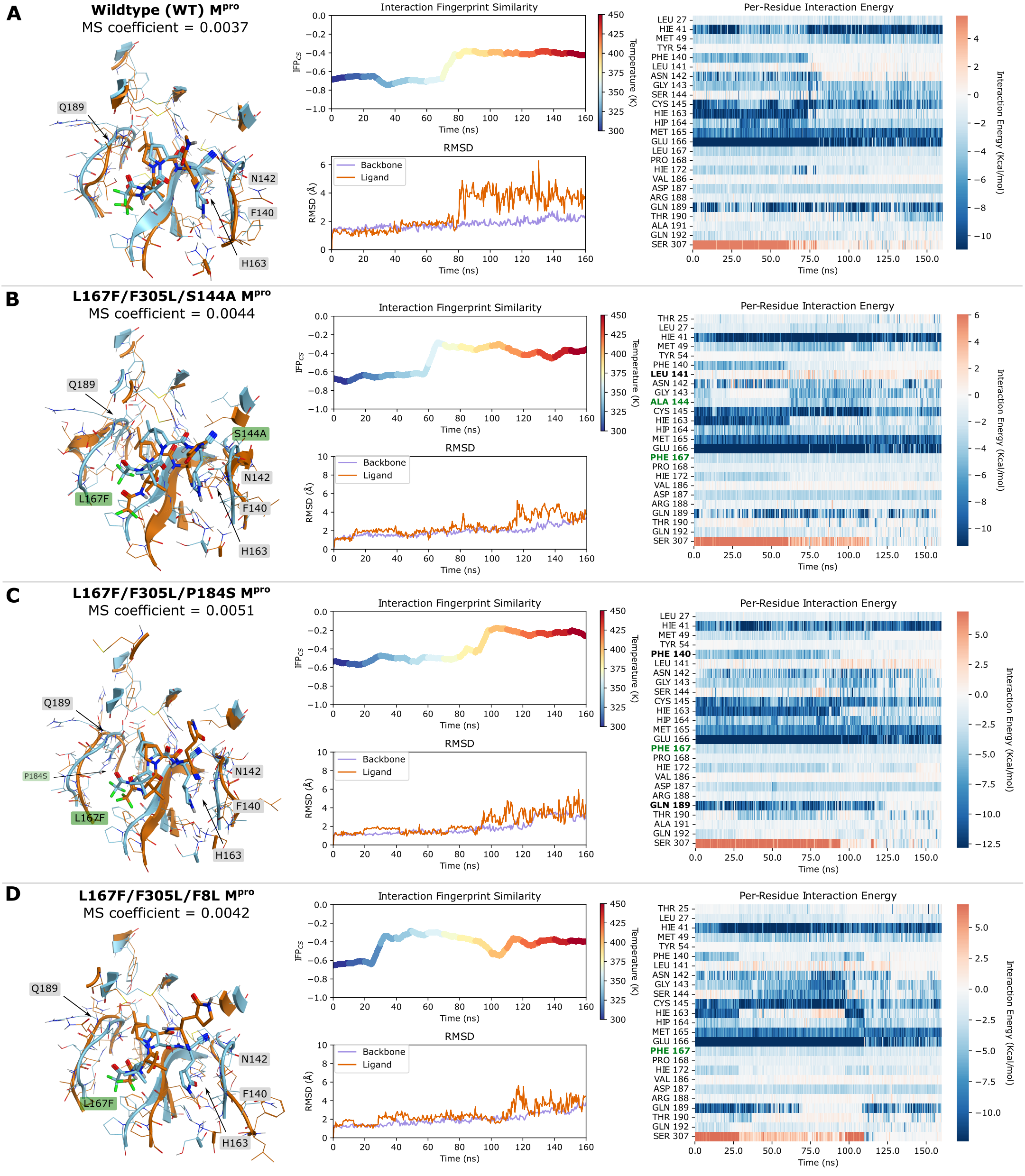

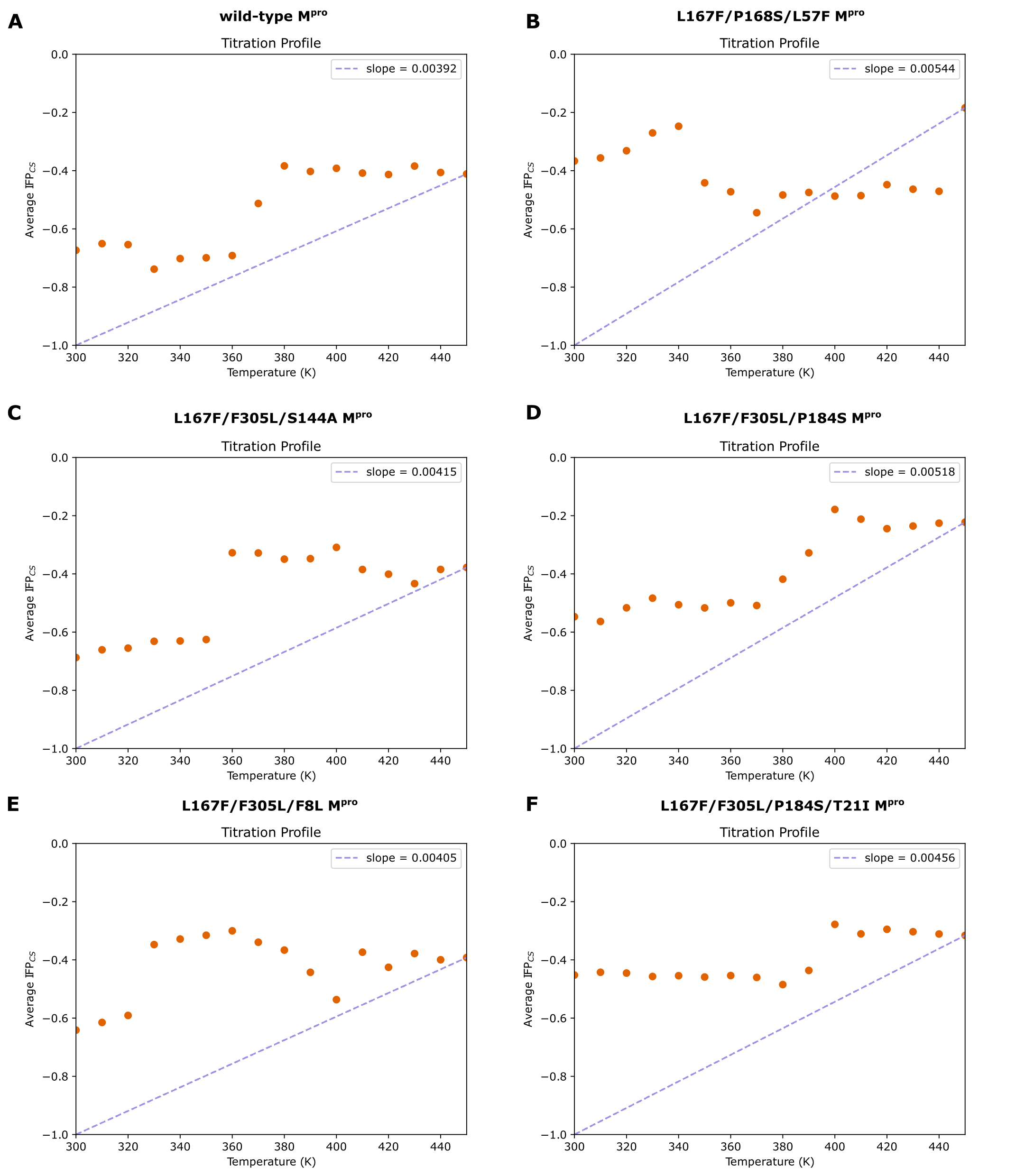

**Fig. S12. MS coefficient plots.** Titration profiles of WT protease and mutants. (**A**) Titration profile of WT M^pro^. MS coefficient = 0.00392. (**B**) Titration profile of L167F/P168S/L57F M^pro^. MS coefficient = 0.00544. (**C**) Titration profile of L167F/F305L/S144A M^pro^. MS coefficient = 0.00415. (**D**) Titration profile of L167F/F305L/P184S M^pro^. MS coefficient 0.00518. (**E**) Titration profile of L167F/F305L/F8L M^pro^. MS coefficient = 0.00405. (**F**) Titration profile of L167F/F305L/P184S/T21I M^pro^. MS coefficient = 0.00456.

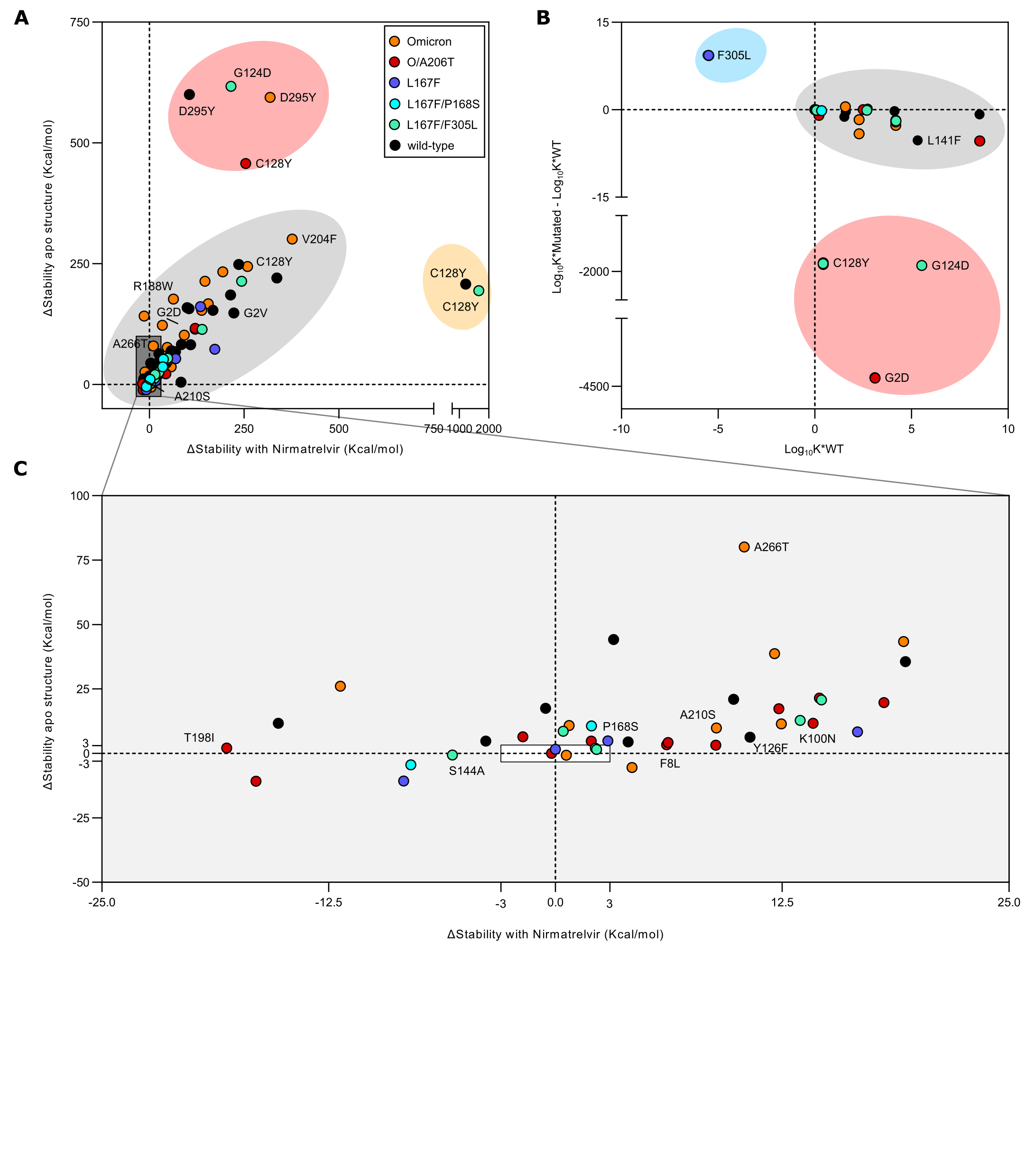

**Fig. S13. Dimerization affinity and stability of M^pro^ apo or nirmatrelvir-bound structures.** Each individual data point represents a substitution and it is coloured according to the parental protease they have been generated from. WT = black; L167F = blue; L167F/F305L = sea green; L167F/P168S = turquoise; Omicron = orange; Omicron/A206T = Bordeaux. (**A**) Plotting of Δ_Stability values for mutations introduced to the apo structure (PDB entry 7ALI) and in the nirmatrelvir bound structure (PDB entry 8DZ2). Coloured ovals represent different groups of mutations based on their destabilization/stabilization of the protease. Red and yellow ovals represent mutants that are strongly off-scale compared to the majority of the other mutants. The light grey ovals represent the third group of mutations that span from slightly stabilizing to strong destabilization. The square coloured in dark grey represents the group of mutations that has closer values to the reference (x=0, y=0). This square is magnified in panel c. (**B**) Dimerization affinity plot. The red oval represents strongly destabilising mutations; the grey oval represents mutations that are borderline-destabilising. The blue oval represents mutations that increase dimerization affinity, in this case F305L only. (**C**) Magnified view of substitutions that cause the least changes in stability of M^pro^. The light grey area around the white box with coordinates (x_1_ = -3, x_2_ = 3; y_1_ = -3, y_2_ = 3) indicates the area where changes in stability are not substantially different. Values outside of these coordinates indicate a substantial change in stability.

| Name | Sequence (5’-3’ direction) |
| --- | --- |
| 33n-before-KpnI-for | GAACCGGTCCTGCTTTCACC |
| G-cut1-rev | CATTTTTCTAAAACCACTCTGCAAAACAGCTGAGGTGATCTTTCCAAGTCGGTTC |
| cut1-for | ATCACCTCAGCTGTTTTGCAG |
| cut2-L-rev | GTCGGTCTCAAAATCGTGGACTTCCATGATTGTTCTTTTCACTGCACTTTG |
| cut2-L-for | AGTGCAGTGAAAAGAACAATCATGGAAGTCCACGATTTTGAG |
| 33n-after-HpaI-rev | GATGTTGGGATGGGATTGGC |
| Omicron-for | CAATGTGCTATGAGGCACAATTTCAC |
| Omicron-rev | CTTAATAGTGAAATTGTGCCTCATAGC |
| blasticidin-for | CATTCGATTAGTGAACGGATCTC |
| L rev | GATGTTGGGATGGGATTGGC |
| hygro-P-for | CTGTTTTGACCTCCATAGAAGATTCTAGAGCTAGCATGGATAATCTCACAAAAGTTC |
| P-hygro-rev | GAGGGAGAGGGGCGGATCCCCTTAATTAACTACAGAGAATATTTGACTCTCGC |
| 3CL^pro^-L167F-for | GCACCATATGGAATTTCCAACTG |
| 3CL^pro^-L167F-rev | CATGAACTCCAGTTGGAAATTCC |
| A206T for | CTATTACAGTTAATGTTTTAACTTGGTTGTACGCT |
| A206T rev | CATTTATAACAGCAGCGTACAACCAAGTTAAAACATT |
| R188W for | GGACCTTTTGTTGACTGGCAAAC |
| R188W rev | CTTGTGCTGTTTGCCAGTCAAC |
| R222L for | GGTGGTTTCTCAATCTATTTACCACAAC |
| R222L rev | GTCATTAAGAGTTGTGGTAAATAGATTGAG |
| Y54H for | CATGCTTAACCCTAATCATGAAGATTTACTC |
| Y54H rev | GACTTACGAATGAGTAAATCTTCATGATTAGGG |
| K100N for | GCCAATCCTAAGACACCTAATTATAAG |
| K100N rev | GCGAACAAACTTATAATTAGGTGTC |
| T198I for | CAGCTGGTACGGACATAAC |
| T198I rev | CATTAACTGTAATAGTTATGTCCGTACC |
| A210S for | GCTTGGTTGTACTCTGCTG |
| A210S rev | CCATTTATAACAGCAGAGTACAACC |
| A266T for | GCCGTTTTAGATATGTGTACTTCATTAAAAG |
| A266T rev | GCAGTAATTCTTTTAATGAAGTACACATATC |
| L167F, P168S for | GCACCATATGGAATTCTCAACTG |
| L167F, P168S rev | CATGAACTCCAGTTGAGAATTCC |
| F8L for | GTTTTAGAAAAATGGCATTACCATCTGG |
| F8L rev | CTCAACTTTACCAGATGGTAATGCC |
| L57F for | CCCTAATTATGAAGATTTCCTCATTCG |
| L57F rev | GATTAGACTTACGAATGAGGAAATCTTC |
| P184S for | GAAGGTAACTTTTATGGATCTTTTGTTG |
| P184S rev | GCCTGTCAACAAAAGATCCATAAAAG |
| S144A for | CATTCCTTAATGGTGCATGTGG |
| S144A rev | CACTACCACATGCACCATTAAG |
| P168S for | CACCATATGGAATTATCAACTGGAG |
| P168S rev | GCATGAACTCCAGTTGATAATTCC |
| T21I for | GGGTTGTATGGTACAAGTAATTTGTGGTAC |
| T21I rev | CGTTAAGTGTAGTTGTACCACAAATTACTTG |
| C145A for | GTTCATTCCTTAATGGTTCAGCTGGTAGTG |
| C145A rev | CTATGTTAAAACCAACACTACCAGCTGAACC |

**Table S1**: cloning primers.

| Residue number | VSV-M^pro^ variant (Nanopore sequencing) | | | | | | | | | |
| --- | --- | --- | --- | --- | --- | --- | --- | --- | --- | --- |
|  | VSV-O-A206T | | VSV-Omicron | | VSV-L167F | | VSV-L167F/P168S | | VSV-L167F/F305L | |
| -4 |  |  |  |  | A(-4)A | 7.50% |  |  |  |  |
| 2 | G2D | 11.89% | G2D | 69.29% |  |  |  |  |  |  |
| 3 | F3S | 22.22% |  |  |  |  |  |  |  |  |
| 8 | F8L | 16.76% |  |  |  |  |  |  | F8L | 8.60% |
| 12 |  |  | K12E | 11.88% | K12K | 7.20% |  |  |  |  |
| 13 |  |  |  |  |  |  |  |  | V13A | 11.66% |
| 17 |  |  |  |  |  |  |  |  | M17L | 5.13% |
| 22 | C22T | 21.41% |  |  | C22G | 37.40% |  |  |  |  |
| 39 |  |  |  |  | P39P | / |  |  |  |  |
| 54 |  |  | Y54H | 28.28% |  |  |  |  |  |  |
| 54 |  |  | Y54H | 25.67% |  |  |  |  |  |  |
| 54 |  |  | Y54H | 12.00% |  |  |  |  |  |  |
| 54 |  |  | Y54H | 6.80% |  |  |  |  |  |  |
| 54 |  |  | Y54H | 6.66% |  |  |  |  |  |  |
| 54 |  |  | Y54H | 4.53% |  |  |  |  |  |  |
| 54 |  |  | Y54H | 15.51% |  |  |  |  |  |  |
| 54 |  |  | Y54H | 50.83% |  |  |  |  |  |  |
| 54 |  |  | Y54H | 12.64% |  |  |  |  |  |  |
| 54 |  |  | Y54H | 13.73% |  |  |  |  |  |  |
| 57 |  |  |  |  |  |  | L57F | 82.27% |  |  |
| 74 |  |  |  |  |  |  | Q74L | 7.68% |  |  |
| 93 |  |  |  |  | T93P | 25.39% |  |  |  |  |
| 98 |  |  |  |  |  |  |  |  | T98S | 4.70% |
| 100 | K100N | 9.11% |  |  | K100K | / |  |  |  |  |
| 100 |  |  |  |  |  |  |  |  |  |  |
| 111 |  |  |  |  |  |  |  |  | T111N | 6.71% |
| 119 | N119D | 11.44% |  |  |  |  |  |  |  |  |
| 119 | N119D | 6.01% |  |  |  |  |  |  |  |  |
| 119 | N119D | 7.40% |  |  |  |  |  |  |  |  |
| 123 | S123S | 10.77% |  |  |  |  |  |  |  |  |
| 124 |  |  |  |  |  |  |  |  | G124D | 6.84% |
| 126 | Y126S | 12.01% |  |  |  |  |  |  |  |  |
| 128 | C128Y | 42.15% | C128Y | 82.48% |  |  |  |  | C128Y | 9.91% |
| 129 | A129S | 9.80% |  |  |  |  |  |  | A129T | 7.26% |
| 139 |  |  |  |  | S139P | 31.30% |  |  |  |  |
| 144 |  |  |  |  |  |  |  |  | S144A | 4.62% |
| 153 | D153N | 6.02% |  |  |  |  |  |  |  |  |
| 168 |  |  |  |  | P168S | / |  |  |  |  |
| 181 |  |  | F181S | 64.62% |  |  |  |  |  |  |
| 184 |  |  |  |  |  |  |  |  | P184S | 21% |
| 188 |  |  | R188W | 72.94% |  |  |  |  |  |  |
| 190 |  |  |  |  | T190T | 12.29% |  |  |  |  |
| 197 |  |  |  |  |  |  |  |  | D197A | 5.13% |
| 198 | T198I | 10.56% |  |  |  |  |  |  |  |  |
| 200 | I200V | 12.31% |  |  |  |  |  |  |  |  |
| 204 |  |  | V204F | 63.18% |  |  |  |  |  |  |
| 206 |  |  | A206T | 94.22% |  |  |  |  |  |  |
| 206 |  |  | A206T | 94.83% |  |  |  |  |  |  |
| 206 |  |  | A206T | 68.01% |  |  |  |  |  |  |
| 206 |  |  | A206T | 37.79% |  |  |  |  |  |  |
| 207 |  |  | W207R | 93.36% |  |  |  |  |  |  |
| 209 |  |  |  |  | Y209Y | 11.31% |  |  |  |  |
| 210 |  |  | A210D | 93.53% |  |  |  |  |  |  |
| 210 |  |  | A210S | 95.15% |  |  |  |  |  |  |
| 210 |  |  | A210T | 38.72% |  |  |  |  |  |  |
| 210 |  |  | A210T | 62.68% | A210T | 10.33% |  |  |  |  |
| 210 |  |  | A210T | 17.96% | A210T | 10.60% |  |  |  |  |
| 210 |  |  |  |  |  |  |  |  |  |  |
| 210 |  |  |  |  |  |  |  |  |  |  |
| 216 |  |  | D216Y | 90.72% |  |  |  |  |  |  |
| 216 |  |  | D216A | 21.02% |  |  |  |  |  |  |
| 220 |  |  | L220P | 76.98% |  |  |  |  |  |  |
| 229 |  |  |  |  | D229G | 13.80% |  |  |  |  |
| 232 |  |  |  |  | L232R | 11.39% |  |  |  |  |
| 234 |  |  | A234D | 7.50% |  |  | A234T | 70.37% |  |  |
| 234 |  |  |  |  |  |  |  |  |  |  |
| 257 | T257N | 15.46% |  |  |  |  |  |  |  |  |
| 260 | A260D | 6.80% |  |  |  |  |  |  |  |  |
| 260 | A260D | 10.88% |  |  |  |  |  |  |  |  |
| 266 |  |  | A266T | 6.60% |  |  |  |  |  |  |
| 268 |  |  |  |  |  |  |  |  | L268F | 55.80% |
| 268 |  |  |  |  |  |  |  |  | L268F | 25.57% |
| 276 | M276R | 7.20% |  |  |  |  |  |  |  |  |
| 277 | N277K | 12.02% |  |  |  |  |  |  |  |  |
| 281 |  |  | I281R | 38.85% |  |  |  |  |  |  |
| 282 |  |  |  |  |  |  | L282S | 7.00% |  |  |
| 289 |  |  | D289T | 49.75% |  |  |  |  |  |  |
| 289 |  |  | D289T | 22.03% |  |  |  |  |  |  |
| 292 |  |  | T292K | 61.08% |  |  |  |  |  |  |
| 295 |  |  | D295V | 60.45% |  |  |  |  |  |  |
| 296 |  |  |  |  |  |  |  |  | V296V | 9.93% |
| 296 |  |  |  |  |  |  |  |  | V296F | 66% |
| 299 |  |  | Q299R | 55.62% |  |  |  |  | Q299R | 23% |
| 305 |  |  |  |  | F305L | / |  |  |  |  |
| (+3) |  |  |  |  | V(+3)V | 9.60% |  |  |  |  |
| (+4) |  |  |  |  | K(+4)T | 9.58% |  |  |  |  |

**Table S2. Retrieved mutations from selection experiments and Nanopore sequencing coverage in percentage (%)**. Table displaying all the retrieved mutations from different VSV-M^pro^ variants used: VSV-O/A206T-M^pro^, VSV-Omicron-M^pro^, VSV-L167F-M^pro^, VSV-L167F/P168S-M^pro^ and VSV-L167F/F305L-M^pro^. For each substitution, the coverage (in percentage) is reported on its right.
